# scResolve: Recovering single cell expression profiles from multi-cellular spatial transcriptomics

**DOI:** 10.1101/2023.12.18.572269

**Authors:** Hao Chen, Young Je Lee, Jose A. Ovando, Lorena Rosas, Mauricio Rojas, Ana L. Mora, Ziv Bar-Joseph, Jose Lugo-Martinez

**Author notes:** These authors contributed equally (Hao Chen, Young Je Lee).

## Abstract

Many popular spatial transcriptomics techniques lack single-cell resolution. Instead, these methods measure the collective gene expression for each location from a mixture of cells, potentially containing multiple cell types. Here, we developed scResolve, a method for recovering single-cell expression profiles from spatial transcriptomics measurements at multi-cellular resolution. scResolve accurately restores expression profiles of individual cells at their locations, which is unattainable from cell type deconvolution. Applications of scResolve on human breast cancer data and human lung disease data demonstrate that scResolve enables cell type-specific differential gene expression analysis between different tissue contexts and accurate identification of rare cell populations. The spatially resolved cellular-level expression profiles obtained through scResolve facilitate more flexible and precise spatial analysis that complements raw multi-cellular level analysis.

## Introduction

The technique of spatial transcriptomics facilitates studying cells *in situ* by combining imaging with the ability of measuring the level of gene expression in different locations of tissue samples. Despite the increasing number of spatial transcriptomics technologies being developed to achieve subcellular-resolution measurements of gene expression [1, 2, 3], the most popular platforms with the ability to sequence the whole transcriptome, such as Spatial Transcriptomics (ST) [4] and Visium [5], still lack the resolution required for single cell level analysis. In such technologies, each measurement spot covers a tissue area of 10 to 20 cells depending on the tissue, a resolution that may be sufficient for well-represented large cells but problematic for small cells or ones that aren’t well represented [6]. Such multi-cellular resolution measurements make it hard to analyze the cell-type organization in tissues and the interactions between cell types or even individual cells.

To overcome resolution limitations, a typical approach involves the use of cell type deconvolution on spatial transcriptomics data. Many methods [7, 8, 9] have been proposed to estimate the composition of cell types in each spot using reference single cell RNA-seq (scRNA-seq) data in which cell types are annotated. Similarly, there are some methods that can perform cell type deconvolution without using the reference scRNA-seq data [10]. More recent methods couple the estimation of gene expression profiles for different cell types with inferring the cell type compositions [10, 11]. However, these methods ignore the heterogeneity of cells within each cell type, that is, even cells originating from the same cell type can exhibit significant variations in their genotypes or phenotypes, influenced by the specific tissue context in which they reside [12]. For example, T cells exhibit an exhausted phenotype in tissues impacted by chronic infections, induced by repeated low-dose stimulation [13], which is different from their normal activation state. Current cell type deconvolution methods do not allow the study of how a certain cell type was developed in response to changes in the tissue environment and to identify the pathways that undergo regulation during these changes. C-SIDE [14] was designed to study the differentially expressed genes in a specific cell type across different tissue contexts. It combines cell type deconvolution with a parametric approach for identifying differential expression. However, it does not compare distinct yet similar cell types. In addition, it relies on predefined cell types in the reference single cell data, restricting its applicability in analyzing newly identified cell types in tissues. On the other hand, computational methods have been proposed to enhance the resolution of multi-cellular spatial transcriptomics data [15, 16], however, these methods are still not appropriate for studying individual cells. Some methods were proposed to reconstruct spatial organizations of single cells by mapping single cells to spatial transcriptomics [17, 18, 19], however, such methods could introduce biases without accounting the batch variation between the single cell dataset and the spatial transcriptomics sections. Crucially, in many cases, access to paired single cell data and spatial transcriptomics data is unavailable.

In this work, we propose a new computational method, *scResolve*, a method for reconstructing the expression profile of each individual cell from spatial transcriptomics data acquired at the multi-cellular resolution. scResolve enables more flexible and detailed spatial analysis at single-cell resolution. Our approach first generates subcellular resolution gene maps by combining spot-level expression profiles with the paired histology image, and then from these maps segments individual cells and thereby produces their expression profiles. We tested scResolve on simulated and real spatial transcriptomics data. As we show, scResolve successfully recovers expression profiles of individual cells in different cell types under different tissue contexts improving upon prior methods. In addition, we demonstrate that cell populations recovered by scResolve are correctly mapped spatially which enables the identification of rare cell populations.

## Results

### Overview of scResolve

scResolve extends SCS [20], which we developed previously for cell segmentation of spatial transcriptomics at subcellular resolution, to the multi-cellular resolution data. SCS performs segmentation in three key steps. It first identifies cell nuclei from tissue staining images. Then a deep learning transformer model is trained to infer for each subcellular spot from gene expression whether it is part of a cell or part of the extracellular matrix, and its relative position with respect to the center of its nucleus. Finally, spots that are determined to be parts of the cells are grouped according to their relative positions to nucleus centers. To enable cell segmentation on the multi-cellular resolution data, we combined SCS with a recently developed super-resolution method, XFuse [16], that integrates spatial gene expression data with histological image data to infer gene expression levels at each pixel in the histology image.

In summary, scResove contains two steps to recover single-cell expression profiles (Fig. 1a). It first enhances the resolution of multi-cellular spatial transcriptomics measurements to pixel resolution, which is at the subcellular level. It next uses SCS to perform cell segmentation on the subcellular resolution gene expression maps. The gene expression values assigned to pixels are ultimately aggregated based on segmentation cell boundaries, resulting in the generation of spatially resolved single-cell expression profiles, which enables flexible single-cell resolution analyses, including cell type organizations in the tissue, cell type-specific differential gene expression, rare cell type identification, and differential gene expression between spatially adjacent cell types (Fig. 1b). See Methods for more details.

**Figure 1:**
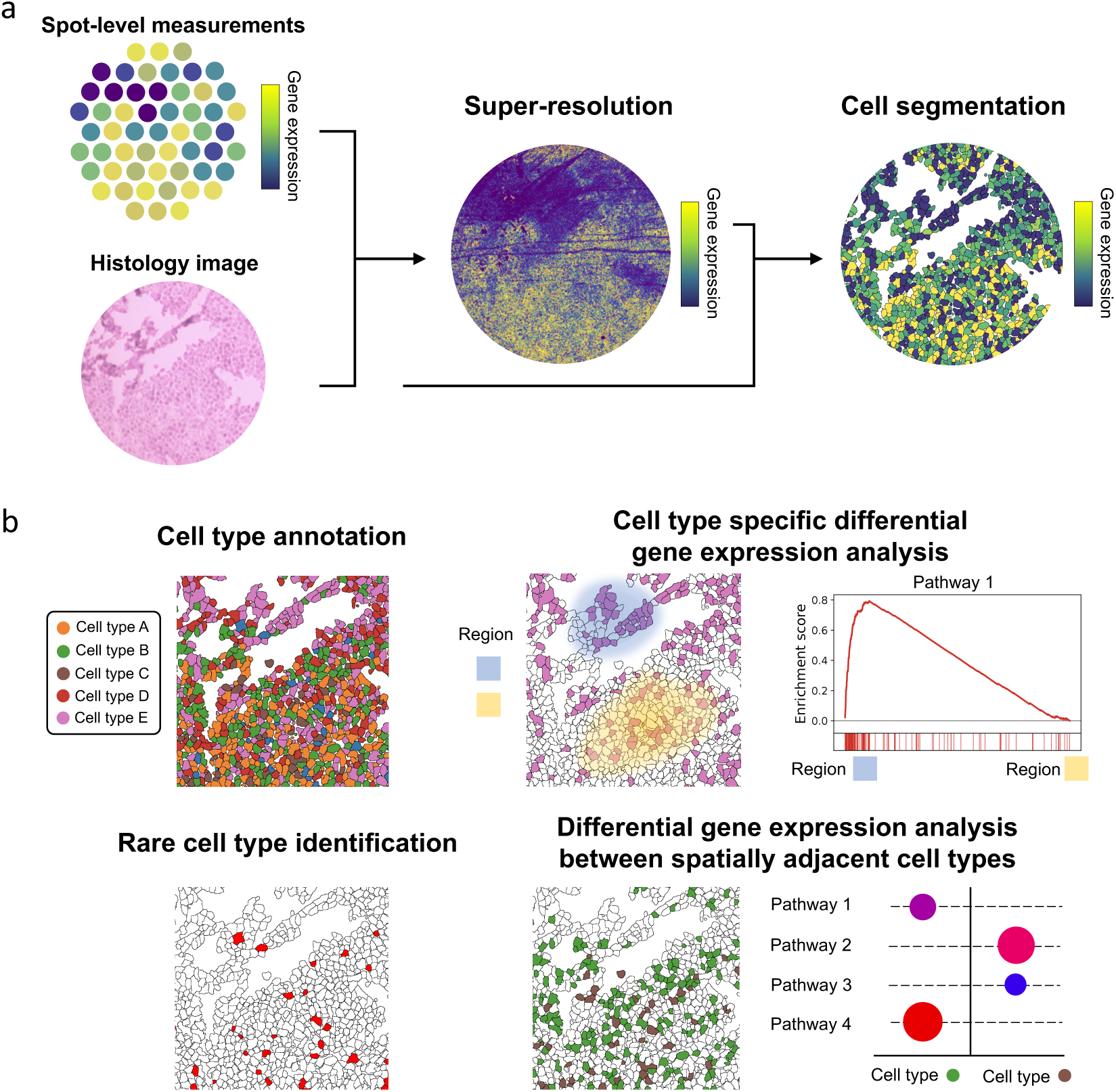
Workflow of scResolve. **a**, scResove contains two steps to recover single-cell expression profiles. It first enhances the resolution of gene expression measurements by inferring the expression levels at each pixel in the histology image using the XFuse model, which takes as input the spot-level gene expression measurements and the histology image. It next uses SCS to perform cell segmentation taking as input the histology image and the inferred subcellular-resolution gene expression maps. The gene expression values assigned to pixels are then aggregated based on the segmentation cell boundaries, resulting in the generation of spatially resolved single-cell expression profiles. **b**, With the expression profile of each recovered single cell in the spatial map, cell type labels can be assigned to cells. Then for each specific cell type, differential gene expression analysis can be performed for cells of that cell type between different tissue contexts. Rare cell types and their locations could be identified. Additionally, the recovered single cell map makes it possible to study the subtle distinctions between cell types that are functionally similar and located in close spatial proximity.

### Evaluation of scResolve and benchmarking on cell type deconvolution

We first evaluated how accurate scResolve is in recovering expression profiles of single cells. For this, we simulated low-resolution spatial transcriptomics data with ground-truth expression profiles of each single cell. Specifically, we randomly placed single cells into a spatial map and divided the map into a grid to simulate low-resolution spots. A corresponding image of the spatial map was generated, where distinct cell types were represented by unique colored shapes (Methods). This simulation enables us to control the distribution of cells across different cell types, allowing for the simulation of rare cell types and providing groundtruth expression profiles for individual cells.

We first applied scResolve to the simulated data section where the numbers of cells for different cell types are evenly distributed (Fig. 2a). From the section of 500 cells, scResolve recovered 491 cells at their original locations, among which 467 cells (95.11%) were then correctly assigned with their respective cell types (Fig. 2b, Methods). This accuracy is significantly higher than that of randomly assigning cell types to the recovered cells (14.29%, *P* -value*<*1.0e-4). To further assess how accurate the recovered gene expression profiles are, for each cell type, we computed the correlation between the average expression profiles of recovered cells assigned to the cell type and actual cells belonging to that cell type. An average Pearson correlation coefficient of 0.98 was obtained across all cell types (Fig. 2c). For the expression correlation between different cell types, the Pearson correlation coefficients obtained between the recovered cells and actual cells are very similar to those obtained between real cells themselves (Figs. 2c-d). With the cell types assigned to our recovered cells, we calculated the cell type compositions in spots, that is, the proportion of different cell types in each spot, and benchmarked them against those derived from the popular cell type deconvolution methods, including Stereoscope[7], cell2location[9], and CARD[8] (Figs. 2e-g, Supplementary Fig. 1, Methods). The cell type compositions obtained from scResolve achieve an average Mean Squared Error (MSE) of 0.066 and an average Pearson correlation coefficient (PCC) of 0.95 with the ground-truth across spots. These values are significantly superior to the results obtained from the other compared methods (average MSE of 0.85 of Stereoscope which is the best among compared methods, *P* -value*<*1.0e-4, average PCC of 0.46 of Stereoscope which is the best among compared methods, *P* -values*<*1.0e-4, Figs. 2h-i). The average performance across five replicates was reported in Supplementary Table 1.

**Figure 2:**
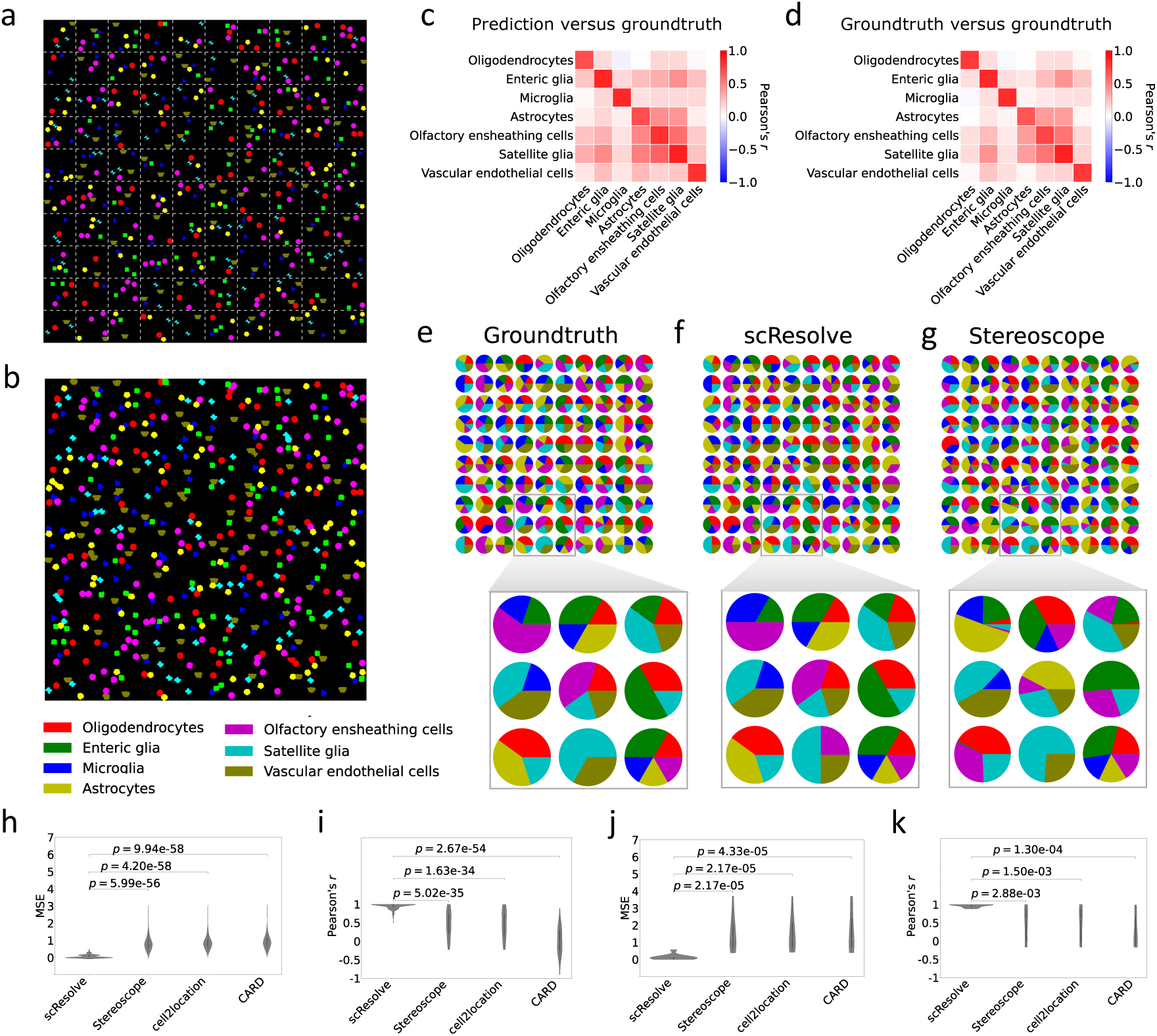
Simulation study. **a**, Image for the synthetic data section with uniformly distributed cell numbers for cell types, where different cell types are represented by unique colored shapes. The whole spatial map was divided into a grid to simulate multicellular spots. **b**, Recovered single cells with their annotated cell types in the spatial map. **c**, Expression correlation between cell types in scResolve recovered cells and those in real cells. **d**, Expression correlation between cell types in real cells. **e**, Groundtruth cell type composition of each simulated spot in **a**. The colors match the colors of cell types in **a**. **f**, Reconstructed cell type compositions from scResolve recovered single cells. **g**, Reconstructed cell type compositions from Stereoscope. **h**, Cell type composition prediction accuracy of different methods for the section in **a**, measured by mean squared error (MSE). Each spot contributes to one point to the errors presented in the violin plot. **i**, Cell type composition prediction accuracy for the section in **a** measured by Pearson correlation coefficient (PCC). **j**, Cell type composition prediction accuracy measured by MSE for a patch of a section simulated with rare cell types. **k**, Cell type composition prediction accuracy measured by PCC for the same patch of the section with rare cell types. One-sided Mann-Whitney U tests were applied between groups.

We then simulated a data section containing two rare cell types where each rare cell type accounts for 2.5% of the population (Supplementary Figs. 2a-b, Methods). scResolve again accurately recovered the expression profiles of the two rare cell types (Supplementary Figs. 2c-d). Remarkably, scResolve accurately recovered the cell type compositions of spots located near the rare cell types (Supplementary Fig. 2a), with an average MSE of 0.13 and an average PCC of 0.97. These values are again significantly better than results obtained for the methods we compared to (average MSE of 1.50 of Stereoscope, *P* -values*<*1.0e-4, average PCC of 0.47 of Stereoscope, *P* -values*<*1.0e-2, Figs. 2j-k). In fact, both Stereoscope and cell2location were only able correctly identify one of the two rare cell types, but failed to recognize the other one (Supplementary Fig. 2e). The average performance across five replicates was again reported in Supplementary Table 1.

### Application of scResolve on human breast cancer dataset

Spatial gene expression adds more detailed information to cancer diagnosis besides analyzing histology images. An important diagnostic issue for breast cancer is to determine the cancer stages, that is, whether the cancer cells have spread or not. Stahl et al. [4] attempted to apply ST technology to human breast cancer tissues to help distinguish stages of cancer. However, due to the low resolution of the data (*∼*50 cells per spot), gene expression can only be studied collectively for multiple cell types in the tumor environment but not for a certain cancer-related cell type individually. This may not influence much on looking at a few marker genes but poses a challenge to systematically study the regulation of pathways in a specific cell type. We applied scResolve to recover single cells from each of the four breast cancer tissue sections in this dataset (Methods). Annotations for areas of invasive ductal cancer (INV, where cancer cells have spread) and areas of ductal cancer in situ (DCIS) were identified on the basis of morphological criteria and are provided in the original publication of this dataset [4] (Fig. 3a, also see Supplementary Fig. 3). This information allows us to assess the prediction results of our method. We first annotated the cell types for the recovered single cells and the original ST spots by label transfer from a reference single cell dataset [21] (Figs. 3b-c, Methods). We found that epithelial cells are enriched in the tumor regions (INV and DCIS regions, 2.73 fold change for single cells, 1.77 fold change for Visium spots), in agreement with the established understanding that breast cancer arises from the breast epithelium [22]. In addition, our recovered single cells revealed the complexity of the tumor environment [23], in which 33.50% recovered cells in the tumor regions are immune cells. However, annotations to the original spots in the tumor regions are homogenous and dominated by epithelial cells, which implies that the information from complex cell types was obscured by that of major cell types in spots.

**Figure 3:**
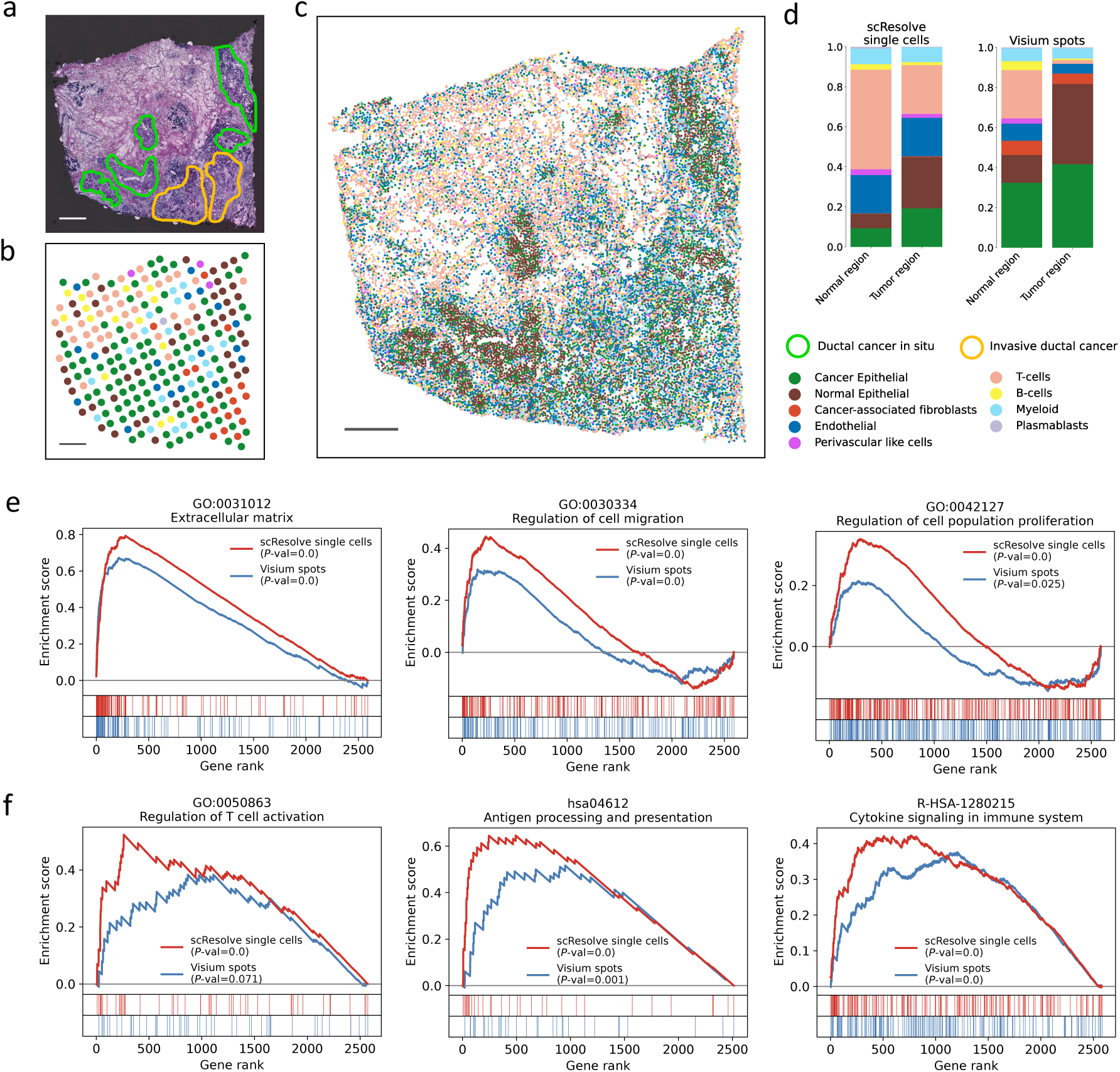
Application to breast cancer tissue. **a**, The histology image of one section in the dataset. Green circles outline ductal carcinoma in situ (DCIS) regions, while yellow circles outline invasive ductal cancer (INV) regions. **b**, Cell type annotations on the original ST spots. **c**, Cell type annotations on the scResolve recovered single cells. **d**, Cell type compositions in different tissue regions derived from the cell type annotations on single cells and spots respectively. **e**, Enrichment of cell migration and proliferation related GO terms in the upregulated genes of epithelial cells in the INV regions. Comparisons were made between differentially expressed genes identified from recovered single cells and those from original spots. **f**, Enrichment of T-cell activation and cytokine signaling related GO terms in the upregulated genes of T-cells in the INV regions. Scale bars, 500 *µ*m.

To further study the heterogeneity of cancer cells in different cancer stages, we performed cell type-specific differential gene expression analysis between the INV and DCIS tumor regions. Specifically, we selected epithelial cells from the INV and DCIS areas respectively, and performed differential gene expression analysis between the two groups of cells. We then looked at the enrichment of gene sets related to cell migration and proliferation, which are known to be active in invasive cancer cells [24, 25], using the GSEA analysis [26] (Methods). The results show that for epithelial cells in INV areas, the upregulated genes are significantly enriched with genes encoding extracellular matrix proteins (*P* -value*<*1.0e-4), as well as those with functions of regulation of cell migration (*P* -value*<*1.0e-4) and regulation of cell population proliferation (*P* -value*<*1.0e-4, Fig. 3e). In comparison, we performed the same analysis for spots between the INV and DCIS regions. Lower enrichment signals were observed for the related gene sets, which suggests the mixture of gene expression of different cell types obscures the differential expression signals for a particular cell type (Fig. 3e). We also looked at T-cells which provide an important line of defense against cancer cells. We performed the same differential gene expression analysis on T-cells and particularly looked at three pathways related to T-cell activation and cytokine signaling [27] using GSEA. We found that compared to T-cells in the other tissue regions, T-cells in INV areas show significant enrichment of the examined pathways in their upregulated genes, suggesting significant tumor-immune interactions in the INV regions. Again, the enrichment signals are all stronger than those identified from the original ST spots (Fig. 3f).

### Application of scResolve on IPF dataset to discover senescent cells

Idiopathic pulmonary fibrosis (IPF) is a progressive and fatal interstitial lung disease for which limited therapeutic options exist [28]. Cellular senescence, a phenomenon characterized by permanent growth arrest and resistance to apoptosis [29] is thought to be involved in IPF pathogenesis [30]. Senescent cells can be initiated by a wide variety of stress-inducing factors [29] and have been shown to induce neighboring normal cells to also undergo senescence through cell-cell contact or by secreting elevated levels of inflammatory cytokines, immune modulators, and growth factors [31]. The accumulation of senescent cells further promotes tissue aging and dysfunction. The first hurdle in removing these detrimental cells involves accurately identifying them in tissues. However, senescent cells are rare in living tissues and so high-resolution spatial data is required to accurately characterize them and their locations. We applied scResolve to four Visium sections of IPF collected from the Senescence Network (SenNet) consortium [32] and identified senescent cells in these sections (Fig. 4a, 4l).

**Figure 4:**
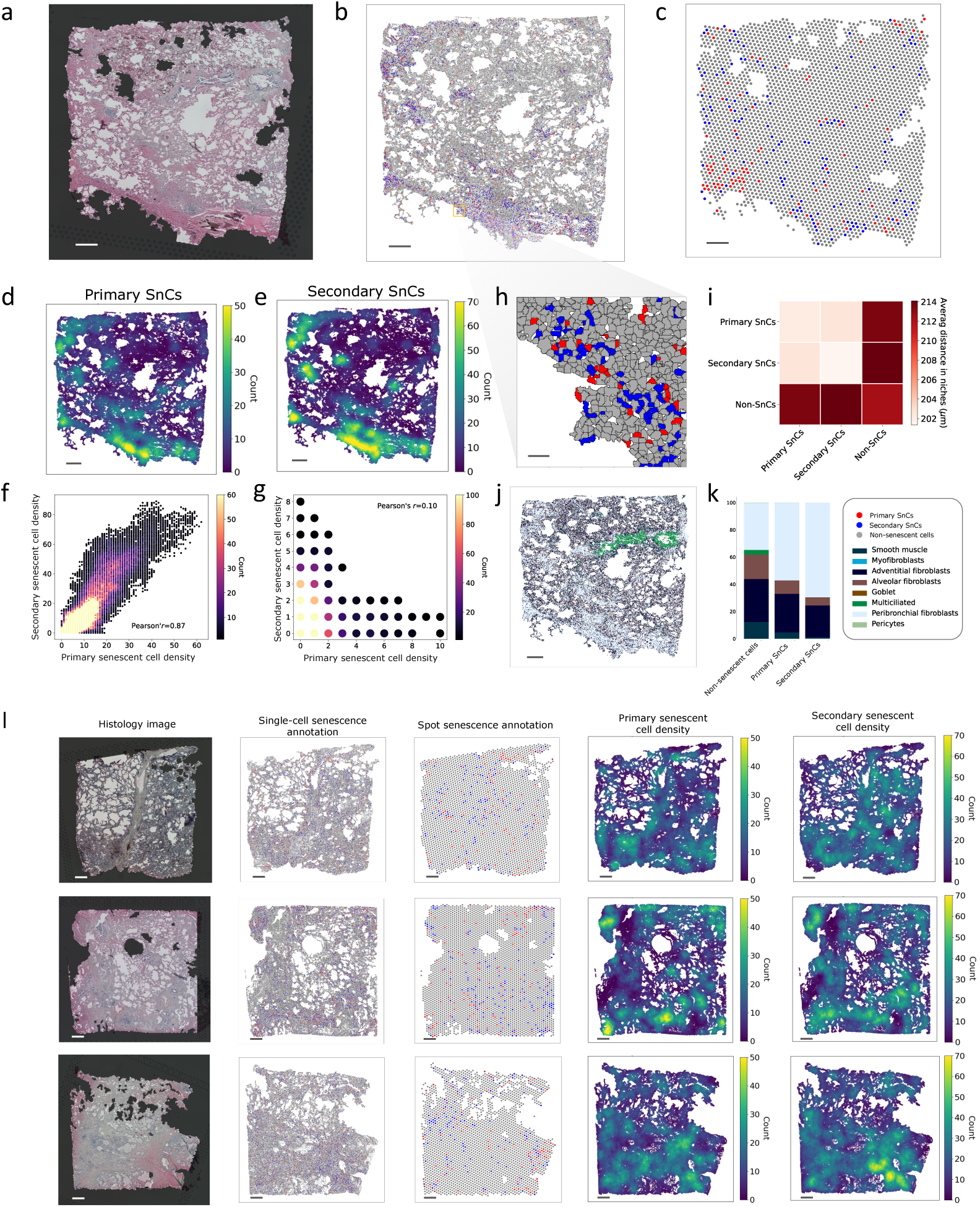
Application to the IPF dataset. **a**, The histology image of one section in the dataset. **b**, Identified primary senescent cells (SnCs), shown as red dots, and secondary senescent cells (blue dots) using the SenMayo gene list in the scResolve-recovered single cells. **c**, Identified spots that contain primary senescent cells (red dots) and secondary senescent cells (blue dots) using the SenMayo gene list. **d**, Primary senescent cell density around each cell in the recovered single cells. **e**, Secondary senescent cell density. **f**, Correlation between the densities of primary and secondary senescent cells. The color of each dot represents the number of cell locations that are surrounded by the densities of senescent cells as indicated by that dot. **g**, Correlation between the densities of primary and secondary senescent cells at the spot level. **h**, Spatial distribution of primary and secondary senescent cells in a small tissue environment (niche). **i**, A heatmap showing the average distance between different groups of cells in niches. **j**, Cell type annotations on the scResolve recovered single cells. **k**, Cell type compositions in different groups of cells. **l**, The results on the other three sections in the dataset. Scale bars, 500 *µ*m in **a-e, j**, 50*µ*m in **h**.

While there are several gene lists of senescent markers [33, 34, 35], significant variations exist between them owing to our limited understanding of senescent cells [36]. This suggests that these lists may capture different aspects of senescent cell features. We first used one of the most recent marker gene lists, SenMayo [33], to annotate the senescent cells in the scResolve recovered cells. Our identified senescent cells account for 3% of all the cells in each of the four sections (Methods). Identified senescent cells are observed to cluster in specific regions within tissue sections (Figs. 4b and 4l). To validate the identified senescent cells, we further sought to identify the population of secondary senescent cells in the tissue sections. The term “secondary” senescent cells is used to distinguish them from intrinsically induced senescent cells (primary senescent cells) by stress factors, such as DNA damage. Secondary senescent cells, on the other hand, are extrinsically induced through interactions with primary senescent cells. Despite having distinct gene expression profiles [37], these two subtypes of senescence should be in close spatial proximity to each other [38]. As suggested, secondary senescent cells were identified using a marker gene list combing the collagen genes and genes in the TGF-*β* signaling pathway [39, 40, 37]. Interestingly, even though the gene lists we used for identifying primary and secondary senescent cells have only 1 gene in common (less than 1% for both lists), the two groups of identified cells show a strong spatial correlation (PCC of 0.89, Figs. 4d-f, Methods), consistent with the established understanding [38]. In contrast, when we applied the same analysis to identify Visium spots containing primary and secondary senescent cells (Figs. 4c and 4l), a much lower spatial correlation was observed between the identified spots for the two types of cells (PCC of 0.10, Fig. 4g, see additional results in Supplementary Fig. 4). In addition, the ability of scResolve to provide single-cell resolution allows for a detailed examination of senescent cell clustering within the tissue microenvironment, also known as niches. We partitioned the entire sections into non-overlapping niches (areas of 1000*×*1000 pixels, approximately 420 *µ*m*×*420 *µ*m). The average distance between different types of senescent cells within niches is smaller compared to the distance between senescent cells and non-senescent cells, as well as between non-senescent cells themselves (Fig. 4i, Supplementary Fig. 5), which reflects the interactions between senescent cells. An example of niches containing primary and secondary senescent cells is shown in Figure 4h. Further cell type annotation on the single cells (Methods) reveals a consistent overrepresentation of peribronchial fibroblasts among senescent cells in four IPF sections (Fig. 4j-k, Supplementary Fig. 6). The proportion of this cell type has been found to increase in IPF [41].

As different gene lists of senescence markers exist with great variations, it is important to test different markers on the same tissues to cross-validate the discoveries. We therefore annotated primary senescent cells using the three other senescence marker lists, FRIDMAN [34], CellAge [35], and Gene Ontology (GO:0090398) [42]. Even though those gene lists do not overlap much (Fig. 5a), the identified senescent cells using different markers show strong spatial correlations (Fig. 5b, Supplementary Figs. 7 and 9). When we tested these gene lists on the original spots, the spatial correlations of identified spots are much lower (Fig. 5c, Supplementary Figs. 8-9). As expected, when we applied these gene lists to the recovered single cells in the sections of healthy lung parenchyma tissues using the same proportion threshold for senescent cells, we generally found weaker clustering patterns of the discovered senescent cells in these sections (Supplementary Fig. 10).

**Figure 5:**
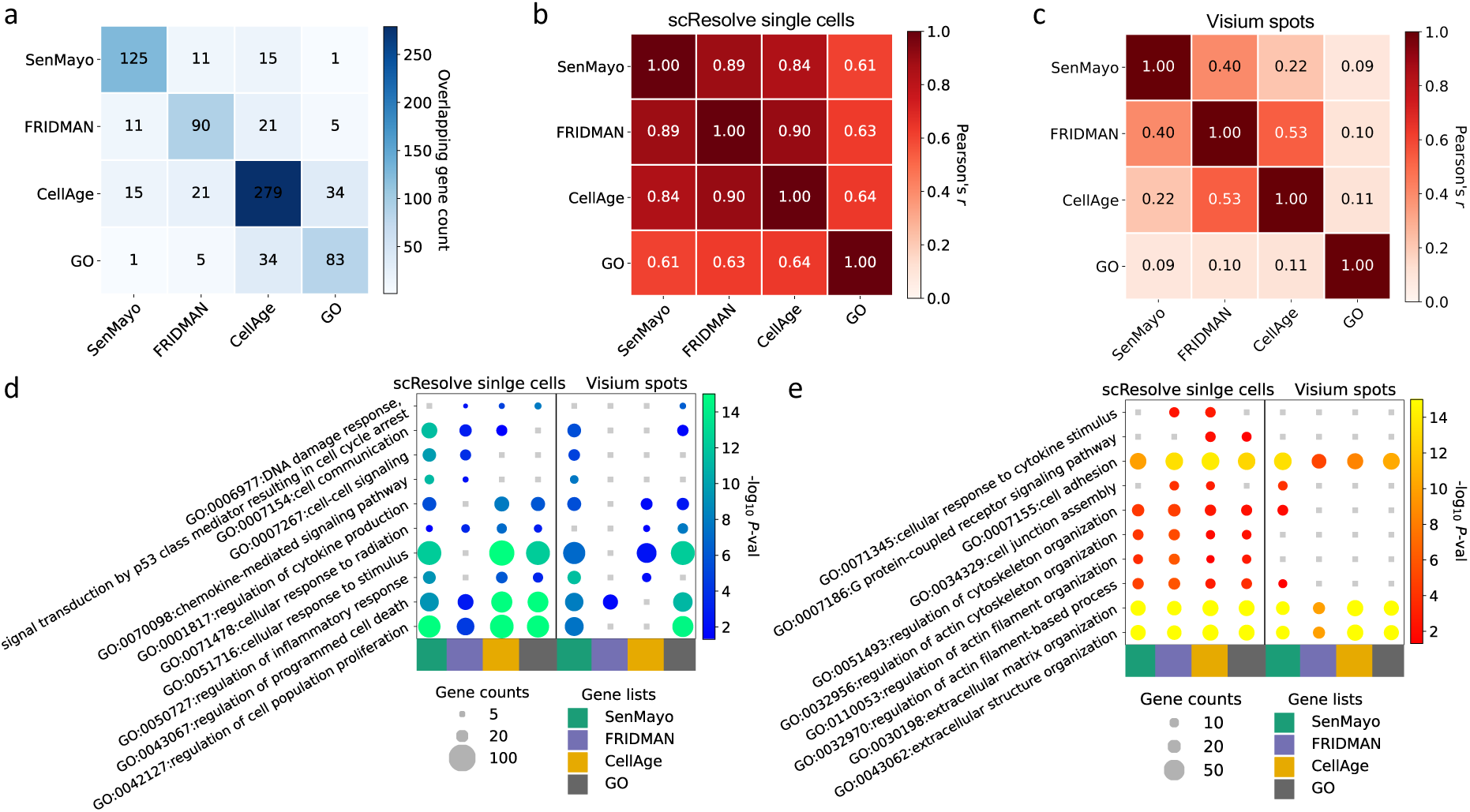
Tests of different senescence makers and differential gene expression analysis between primary and secondary senescent cells. **a**, The count of genes that overlap between different lists of senescent markers. **b**, The spatial correlation among the densities of primary senescent cells identified using different marker gene lists. **c**, The spatial correlation among the densities of spots containing primary senescent cells identified using different marker gene lists. **d**, Enrichment analysis of Gene Ontology (GO) in the top 300 differentially expressed genes in primary senescent cells compared to secondary senescent cells. The size of the dots indicates the number of genes associated with the GO terms present in the differentially expressed genes, while the color of the dots reflects the corresponding *P* -values. **e**, GO enrichment analysis in the top 300 differentially expressed genes in secondary senescent cells.

Finally we performed differential gene expression analysis between identified primary and secondary senescent cells, and repeated the analysis for the senescent cells identified using different marker gene lists. We selected the top 300 upregulated genes for both the primary senescent cells and secondary senescent cells respectively based on the *P* -values from the t-test (Methods). Gene Ontology (GO) enrichment analysis was further performed for the selected genes. GO terms associated with the current understanding of senescent cells and secondary senescent cells are enriched in the respective gene lists of the two cell types. For example, GO terms such as regulation of programmed cell death, and regulation of inflammatory response were found to be enriched for primary senescent cells, in agreement with known pathways for these cells [43, 44] (Fig. 5d). GO terms related to extracellular matrix remodeling were observed for secondary senescent cells, which have been found to be the feature of senescent cells in their early stages [39, 40] (Fig. 5e). When we performed a similar analysis at the spot level, we observed lower enrichment levels for the GO terms related to the known pathways for both primary and secondary senescent cells (Figs. 5d-e). We also performed GO enrichment analysis using the top 300 upregulated genes after excluding marker genes from consideration. Enrichment for most of the GO terms persisted in our single-cell resolution (Supplementary Fig. 11).

## Discussion

The advancement of spatial transcriptomics has enabled the study of cells within their tissue environments. However, many widely used platforms in spatial transcriptomics still only achieve multicellular resolution, where each spot measures the collective expression of multiple cells. This constraint restricts their utility in studying individual cells or specific cell types. To surmount this challenge, we developed scResolve, a computational approach that integrates two deep learning models to reconstruct single-cell expression profiles from low-resolution spatial transcriptomics data. scResolve first predicts subcellular resolution gene expression maps using histology images and multi-cellular gene expression measurements. It then segments individual cells from the resultant high-resolution expression maps. In our simulation study, we demonstrated that scResolve effectively restores the expression profile of each individual cell, closely resembling that of the original cell at its location.

Applications of scResolve on two datasets generated from different multicellular spatial transcriptomics platforms demonstrate the distinct benefits of scResolve-recovered single cells compared to the original multicellular spots. In the human breast cancer dataset, scResolve enables cell type-specific differential gene expression analysis, leading to more significant identifications of regulated pathways within specific cell types across tissue areas with different cancer stages. In addition, scResolve enables the identification of primary and secondary senescent cells at the single-cell resolution in IPF tissue sections. Compared to the spot level, the recovered single cells provide a more accurate identification of senescence. These analyses complement the multi-cellular level analyses that can be performed in the raw data, which usually focus on cell type deconvolution [7, 10] or spatial domains in tissues [45, 46].

While we have demonstrated the efficacy of scResolve on datasets from two different platforms, there is room for further improvement. In the inference of subcellular resolution spatial gene expression maps, each gene is associated with a set of metagenes, each characterizing a distinct expression pattern. The accuracy of gene expression inference is influenced by the number of metagenes. Exploring an algorithm that identifies the minimum number of metagenes without losing information could enhance model robustness. In addition, scResolve currently operates in two separate steps with two distinct models. Designing an end-to-end model could optimize the entire pipeline by consolidating different modules under a unified learning objective, and potentially make it more computationally efficient. To develop more effective methods for this task, evaluating performance on real data with ground-truth is desired. This could be accomplished if there is multi-omics data available, which applies low-resolution spatial transcriptomics and high-resolution spatial proteomics on the same tissue section or adjacent tissue sections.

scResolve is available for download at: https://github.com/chenhcs/scResolve. With the increasing popularity of spatial transcriptomics in cell biology studies[7], diseases pathogenesis analysis [47], and drug discovery[48], we hope scResolve would be a useful tool that enables more precise and flexible spatial analysis of such data.

## Acknowledgements

We thank L.Bergenstråhle and J.Lundeberg for providing insightful responses to our inquiries about XFuse. This work was partially supported by National Institutes of Health (NIH) grants OT2OD026682, and 1U24CA268108 to Z.B.-J., and 1U54AG075931 to M.R., A.L.M., Z.B.-J. and J.L.-M.

## Author Contributions

H.C. and J.L.-M. conceptualized and designed the study. H.C. and Y.J.L. developed the software of scResolve. Y.J.L., H.C., and J.L.-M. performed evaluations and the simulation study. H.C., Y.J.L., and J.L.-M. performed the analyses for the breast cancer dataset. H.C., Y.J.L., J.A.O., L.R., M.R., A.L.M, and J.L.-M. performed the analyses for senescent cells in IPF tissue. H.C., Y.J.L., Z.B.-J., and J.L.-M. wrote the manuscript. All authors read and approved the final manuscript.

## Competing Interests

The authors declare no competing interests.

## Methods

### Super-resolution of multi-cellular spatial expression data

scResolve uses XFuse[1] to improve the resolution of spatial gene expression measurements from the multi-cellular spot level to the granularity of pixels in the histology image. XFuse models the expression level of each gene *g* at each pixel *i* in the histology image as a negative binomial distribution *X_gi_ ∼ NB*(*r_gi_, p_g_*), where the parameters of the distribution, the number of failures before stopping *r_gi_* and the success probability *p_g_*, are mapped from the input histology image *I* through encoder and decoder convolutional neural networks. The pixel-level expression *X_gi_* is related to the gene expression *X_gl_*in each measurement spot *l* by summing over the pixels *A_l_* in the spot, *X_gl_ ≡* ^L^*_i∈A_ X_gi_*. Therefore, the model can be learned from the observational data 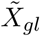 using variational inference.

The parameter *r_gi_*in the distribution of each gene *g* in each pixel *i* is associated with a set of hidden factors, called *metagenes*. The number of metagenes is optimized during model training. The more complex the tissue environment is, typically the more metagenes are needed. In practice, we observed that too few metagenes can lead to inaccurate inference of gene expression. We therefore introduced a user-defined hyperparameter *m*, to retain the latest model checkpoint with at least *m* metagenes. The hyperparameter *m* can be set based on the anticipated number of cell types in the tissue section.

After the model was trained, gene counts were sampled from the learned distributions. For each gene in each pixel, we performed the sampling for 10 times, and the least integer greater than or equal to the mean value of the sampled counts was used. A gene expression map *X* at the pixel level was finally generated. Due to the large amount of computations required by XFuse, we trained a model for each section in each dataset. For each section, we considered only genes with at least 50 counts summed across all the spots, which is recommended by XFuse.

### Cell segmentation from super-resolution spatial expression data

scResolve uses SCS[2] to identify cell boundaries from the super-resolution gene expression map *X*. SCS segments cells by first identifying cell nuclei from the histology image *I* using watershed algorithm[3]. It then uses a transformer model to infer for each pixel *i* the object probability *p_i_ ∈* [0, 1], that is, the probability the pixel is part of a cell versus part of the extracellular matrix, and a two-dimensional gradient 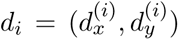, representing the direction from the pixel to the center of its cell nucleus. To make predictions for each pixel, the transformer model adaptively aggregates high-dimensional but sparse gene expression information from neighboring pixels via an attention mechanism. For model training, SCS uses as positive examples pixels within the identified nuclei and as negative examples pixels sampled from highly confident background regions. Finally, pixels with object probabilities greater than a threshold *θ* are determined to be part of the cells and grouped by tracking the gradient flow from pixels to nucleus centers, in which, starting from a pixel *i* = (*x, y*), the next pixel to which the flow is directed is selected using 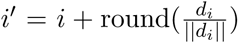. The gene counts of pixels determined to belong to the same cell are summed as the gene expression for that cell.

The watershed algorithm of SCS takes one channel image as input, and detects bright regions as nuclei. scResolve converts the original H&E images into grayscale and inverts the pixel values in the grayscale images to make the nucleus regions brighter than the cytoplasm and extracellular regions. Due to the large size of the tissue sections in our datasets, we cut each section into nonoverlapping patches and run SCS on each patch separately.

### Generation of synthetic data

We generated synthetic multi-cellular spatial data using scRNA-seq data from the Atlas of the Adolescent Mouse Brain[4]. Seven cell types were used for the simulation, including Oligodendrocyte, Enteric glia, Microglia, Astrocyte, Olfactory ensheathing cell, Satellite glia, and Vascular endothelial cell. The cell types were selected based the sum of the scores (*t* statistics) of the top 20 differentially expressed genes of each cell type. The seven cell types with the highest sum of scores were selected to ensure that the selected cell types have larger genotypic differences. Each section in the synthetic data is a spatial map of size 1000*×*1000 pixels, which was divided into 100 spots, each with 100*×*100 pixels in size.

We then placed single cells in the spatial map. The number of cells in each spot was sampled from the normal distribution with mean 5 and variance 1. For the case of uniformly distributed cell types, for each cell to be placed in each spot, we sampled a cell type from the uniform distribution, and randomly picked a cell from the selected cell type and placed it at a random location in the spot. For the case with simulated rare cell types, two cell types were randomly picked as rare cell types. We then sampled cells from a cell type distribution in which each of the two rare cell types accounts for 2.5% of the population and the other five cell types evenly occupy the remaining 95% of the population. Cells of each rare cell type were placed at two adjacent randomly selected spots and cells from the other cell types were randomly distributed across all spots. The gene expression profile of each spot was calculated as the sum of gene counts of cells in the spot. The union of the top 20 differentially expressed genes of each cell type, 117 genes in total, were used to define the gene scope for the simulated data.

We generated an image for the spatial map. In the image, seven cell types were represented with colored shapes of red circles, green squares, blue triangles, yellow hexagons, pink octagons, cyan ribbons, and brown trapezoids, respectively. The shapes of size around 16*×*16 pixels were placed at the corresponding locations of sampled cells in the map with a black background.

For running scResolve on the synthetic data. The hyperparameter *m* for the number of metagenes was set to 7. SCS was applied to the entire section of each simulated spatial map. The threshold *θ* for object probability was set to 0.1 as the default.

### Benchmark comparison with cell type deconvolution methods

To benchmark with existing methods, we first annotated the cell type of each recovered cell of scResolve. The fold change of each gene in each cell was calculated by comparing the expression of the gene in that cell with the average expression of the gene in all cells. Then the 20 genes with the highest fold changes were selected from each cell. The cell type label for each cell was assigned as the cell type whose top 20 differentially expressed genes have the greatest overlap with the 20 selected genes of that cell. To quantify the cell type annotation accuracy, recovered cells were mapped to the original cells in the spatial map image based on their cell mask overlaps with the original cells. After annotating the cell types for cells in each spot, we calculated the cell type composition for each spot. We then compared these compositions with those derived from the popular cell type deconvolution methods, including, Stereoscope[5], cell2location[6], and CARD[7], using the ground-truth cell type compositions as a benchmark.

For each spot *l*, the mean squared error (MSE) between the predicted cell type composition vector 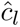 and the ground-truth cell type composition vector *c_l_* was calculated:

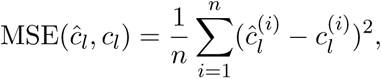

where *n* is the number of cell types. The Pearson correlation coefficient (PCC) between the two vectors 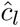 and *c_l_*was also calculated:

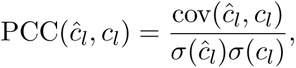

where cov is the covariance, and *σ* is the standard deviation.

For Stereoscope, we used the default parameters, except for using the batch size parameters of 10 for both single cell data and spatial data, to accommodate our simulated data size. We followed the guidance to use cell2location. Instead of using filtered genes, we used all the selected genes in our dataset. For user-specified hyperparameters, we set the estimated number of cells per spot as 5 and tuned the parameter for within-slide variation in RNA detection sensitivity for its best performance. Training epochs for the estimation of reference cell type signatures were set as 3000 and for the spatial mapping of cell types were set as 40,000 to ensure convergence. For CARD, we followed the default setting, except for using parameters minCountGene as 1 and minCountSpot as 1 to consider all the genes and spots.

### Application to the human breast cancer dataset

The human breast cancer spatial transcriptomics dataset was obtained from the original publication [8] (https://www.spatialresearch.org/resources-published-datasets/doi-10-1126science-aaf2403/), which contains four adjacent tissue sections. The data were generated from the Spatial Transcriptomics (ST) platform, where the diameter of each measurement spot is 100 *µ*m and the spot center-to-center distance is 200 *µ*m. The generated data were preprocessed using the ST pipeline (https://github.com/ SpatialTranscriptomicsResearch/st pipeline). For all datasets, spots outside the tissue have been removed.

When running XFuse, after filtering out the genes with at least 50 counts summed across all the spots, there are 3475, 3368, 3190, and 3413 genes kept in four sections respectively, among which 2598 genes are common across all four sections. The size of each H&E image was scaled down to 30% on both axes of the original image to speed up the inference of XFuse, while still maintaining sufficient resolution for cell segmentation. The hyperparameter *m* for the number of metagenes was set to 20. When running SCS, each section was evenly cut into 4*×*4 patches, each with a size of around 2000*×*2000 pixels (940 *µ*m *×* 940 *µ*m). In each patch, 2,000 variable genes were identified as the scope of genes for the expression profiles of spots. The threshold *θ* for object probability was set as 0.1 as the default.

### Application to the human IPF dataset

The spatial transcriptomics dataset of IPF was obtained from SenNet consortium [9]. The data collection protocol and procedure can be found in Nguyen et al. [10]. Four IPF tissue sections from the same donor are contained in this dataset, in which the first two sections are from the upper lobe while the rest two sections are from the lower Lobe. The data were generated from the Visium platform. The diameter of each measurement spot is 65 *µ*m and the spot center-to-center distance is 100 *µ*m. The resulting data were pre-processed using Space Ranger of 10x Genomics. We rely on the tissue mask generated by Space Ranger to filter out spots outside the tissue region.

There are 14,326, 15,536, 14,307, and 12,537 genes kept in four sections respectively after filtering out the genes with at least 50 counts summed across all the spots for running XFuse, among which 12,339 genes are common across all four sections. The resolution of each H&E image was reduced to 30% pixels of the original image. The hyperparameter *m* for the number of metagenes was set to 20. Due to the complexity of the IPF lung tissues, for running SCS, each section was cut into 14*×*14 patches, each with a size of around 1000*×*1000 pixels (420 *µ*m*×*420 *µ*m). In each patch, 2,000 variable genes were identified as the scope of genes for the expression profiles of spots. The threshold *θ* was set as 0.1 as the default.

### Cell type annotation

We transferred cell type labels from the single cell atlas of human breast cancer [11] to the scResolve recovered single cells in the breast cancer dataset. The ingest method implemented in Scanpy [12] was used for label transferring, in which the scRNA-seq dataset in the breast cancer single cell atlas [11] was used as the reference data, while the scResolve recovered single cells were treated as the query data. The intersection of genes between the two datasets was taken to make them defined on the same variables. The ingest data integration function fits a model of PCA combined with a neighbor lookup search tree on the reference data and uses it to project our scResolve recovered single cells. The cell type categories defined at the level of major cell types in the reference single cell data were used. Label transferring was run for each section separately. We used the same method to annotate the original ST spots.

We transferred cell type labels of single cells in the Human Lung Cell Atlas (HLCA) [13] to scResolve recovered single cells in the IPF dataset. We directly used the trained scArches [14] model provided by HLCA for label transfering. scArches performs joint embedding of reference data and query data and uses a nearest neighbor classifier to label cell types. We chose cell types at level 4 among the five levels defined in HLCA, ranging from level 1 (coarsest) to level 5 (most fine-grained).

### Senescent cell identification

To annotate primary senescent cells, we ranked genes within each cell based on their expression values. We then determined for each cell the count of senescence marker genes within its top 10% of genes with the highest expression levels. Cells were then ranked based on these counts, and those in the top 3% were identified as primary senescent cells. For annotating secondary senescent cells, we utilized a gene list derived from the union of collagen genes and genes in the TGF-*β* signaling pathway as markers [15, 16, 17]. The same ranking method was applied, and cells that were ranked in the top 5% and were not identified as primary senescent cells were annotated as secondary senescent cells.

To determine the density of senescent cells surrounding the location of each cell, we counted the number of senescent cells within a distance of 500 pixels (210 *µ*m) from the cell. The spatial correlation between primary and secondary senescent cells was calculated by the Pearson correlation between the densities of the two types of cells across cell locations. The same method was applied to calculate the spatial correlation between primary and secondary senescent cells at the spot resolution.

### Differential gene expression and enrichment analysis

We used t-test to identify genes that are differentially expressed between two groups of cells. Specifically, for the human breast cancer dataset, we first gathered all the epithelial cells, including normal epithelial and cancer epithelial, from the tumor regions across all four sections, and grouped them based on their locations in the areas of ductal cancer in situ (DCIS) or invasive ductal cancer (INV). The t-test was performed between the two groups of cells. The GSEA analysis was next run for the INV group on the preranked genes with their *t* statistics from the t-test. Following this, we selected all the T-cells from the four sections and grouped them based on their locations in the areas of DCIS, INV, or normal tissue. The GSEA analysis was again run for the T-cells in the INV region on the preranked genes with their *t* statistics. For the IPF dataset, a t-test was conducted between primary senescent cells and secondary senescent cells across all four sections. The top 300 differentially expressed genes were identified for each group of cells. GO enrichment was performed in the 300 differentially expressed genes using Fisher’s exact test for primary senescent cells and secondary senescent cells respectively.

## Data availability

The histology images and spatial transcriptomics data for the breast cancer dataset are available from the database of the original publication (https://www.spatialresearch.org/resources-published-datasets/doi-10-1126science-aaf2403/). The histology images and spatial transcriptomics data for the IPF and healthy lung tissue sections were collected within the Cellular Senescence Network (SenNet) Consortium and will be available in the SenNet Data Portal (https://data.sennetconsortium.org/).

## Code availability

The source code of scResolve is publicly available at https://github.com/chenhcs/scResolve.

## Notes

### Competing Interest Statement

The authors have declared no competing interest.

## References

[1] Ao Chen, Sha Liao, Mengnan Cheng, Kailong Ma, Liang Wu, Yiwei Lai, Xiaojie Qiu, Jin Yang, Jiangshan Xu, Shijie Hao, et al. Spatiotemporal transcriptomic atlas of mouse organogenesis using dna nanoball-patterned arrays. Cell, 185(10):1777–1792, 2022.

[2] Chun-Seok Cho, Jingyue Xi, Yichen Si, Sung-Rye Park, Jer-En Hsu, Myungjin Kim, Goo Jun, Hyun Min Kang, and Jun Hee Lee. Microscopic examination of spatial transcriptome using seq-scope. Cell, 184(13):3559–3572, 2021.

[3] Xiaonan Fu, Li Sun, Runze Dong, Jane Y Chen, Runglawan Silakit, Logan F Condon, Yiing Lin, Shin Lin, Richard D Palmiter, and Liangcai Gu. Polony gels enable amplifiable dna stamping and spatial transcriptomics of chronic pain. Cell, 185(24):4621–4633, 2022.

[4] Patrik L Ståhl, Fredrik Salmén, Sanja Vickovic, Anna Lundmark, Jośe Ferńandez Navarro, Jens Magnusson, Stefania Giacomello, Michaela Asp, Jakub O Westholm, Mikael Huss, et al. Visualization and analysis of gene expression in tissue sections by spatial transcriptomics. Science, 353(6294):78–82, 2016.

[5] Lambda Moses and Lior Pachter. Museum of spatial transcriptomics. Nature Methods, 19(5):534–546, 2022.

[6] Cameron G Williams, Hyun Jae Lee, Takahiro Asatsuma, Roser Vento-Tormo, and Ashraful Haque. An introduction to spatial transcriptomics for biomedical research. Genome Medicine, 14(1):1–18, 2022.

[7] Alma Andersson, Joseph Bergenstråhle, Michaela Asp, Ludvig Bergenstråhle, Aleksandra Jurek, Jośe Ferńandez Navarro, and Joakim Lundeberg. Single-cell and spatial transcriptomics enables probabilistic inference of cell type topography. Communications biology, 3(1):565, 2020.

[8] Ying Ma and Xiang Zhou. Spatially informed cell-type deconvolution for spatial transcriptomics. Nature biotechnology, 40(9):1349–1359, 2022.

[9] Vitalii Kleshchevnikov, Artem Shmatko, Emma Dann, Alexander Aivazidis, Hamish W King, Tong Li, Rasa Elmentaite, Artem Lomakin, Veronika Kedlian, Adam Gayoso, et al. Cell2location maps fine-grained cell types in spatial transcriptomics. Nature biotechnology, 40(5):661–671, 2022.

[10] Brendan F Miller, Feiyang Huang, Lyla Atta, Arpan Sahoo, and Jean Fan. Reference-free cell type deconvolution of multi-cellular pixel-resolution spatially resolved transcriptomics data. Nature communications, 13(1):2339, 2022.

[11] Tinyi Chu, Zhong Wang, Dana Pe’er, and Charles G Danko. Cell type and gene expression deconvolution with bayesprism enables bayesian integrative analysis across bulk and single-cell rna sequencing in oncology. Nature Cancer, 3(4):505–517, 2022.

[12] Masoud Najafi, Nasser Hashemi Goradel, Bagher Farhood, Eniseh Salehi, Somaye Solhjoo, Heidar Toolee, Ebrahim Kharazinejad, and Keywan Mortezaee. Tumor microenvironment: Interactions and therapy. Journal of cellular physiology, 234(5):5700–5721, 2019.

[13] Alex D Waldman, Jill M Fritz, and Michael J Lenardo. A guide to cancer immunotherapy: from t cell basic science to clinical practice. Nature Reviews Immunology, 20(11):651–668, 2020.

[14] Dylan M Cable, Evan Murray, Vignesh Shanmugam, Simon Zhang, Luli S Zou, Michael Diao, Haiqi Chen, Evan Z Macosko, Rafael A Irizarry, and Fei Chen. Cell type-specific inference of differential expression in spatial transcriptomics. Nature methods, 19(9):1076–1087, 2022.

[15] Edward Zhao, Matthew R Stone, Xing Ren, Jamie Guenthoer, Kimberly S Smythe, Thomas Pulliam, Stephen R Williams, Cedric R Uytingco, Sarah EB Taylor, Paul Nghiem, et al. Spatial transcriptomics at subspot resolution with bayesspace. Nature biotechnology, 39(11):1375–1384, 2021.

[16] Ludvig Bergenstråhle, Bryan He, Joseph Bergenstråhle, Xesús Abalo, Reza Mirzazadeh, Kim Thrane, Andrew L Ji, Alma Andersson, Ludvig Larsson, Nathalie Stakenborg, et al. Super-resolved spatial transcriptomics by deep data fusion. Nature biotechnology, 40(4):476–479, 2022.

[17] Milad R Vahid, Erin L Brown, Chlóe B Steen, Wubing Zhang, Hyun Soo Jeon, Minji Kang, Andrew J Gentles, and Aaron M Newman. High-resolution alignment of single-cell and spatial transcriptomes with cytospace. Nature Biotechnology, pages 1–6, 2023.

[18] Qihuang Zhang, Shunzhou Jiang, Amelia Schroeder, Jian Hu, Kejie Li, Baohong Zhang, David Dai, Edward B Lee, Rui Xiao, and Mingyao Li. Leveraging spatial transcriptomics data to recover cell locations in single-cell rna-seq with celery. Nature communications, 14(1):4050, 2023.

[19] Jingyang Qian, Jie Liao, Ziqi Liu, Ying Chi, Yin Fang, Yanrong Zheng, Xin Shao, Bingqi Liu, Yongjin Cui, Wenbo Guo, et al. Reconstruction of the cell pseudo-space from single-cell rna sequencing data with scspace. Nature Communications, 14(1):2484, 2023.

[20] Hao Chen, Dongshunyi Li, and Ziv Bar-Joseph. Scs: cell segmentation for high-resolution spatial transcriptomics. Nature methods, 20:1237–1243, 2023.

[21] Sunny Z Wu, Ghamdan Al-Eryani, Daniel Lee Roden, Simon Junankar, Kate Harvey, Alma Andersson, Aatish Thennavan, Chenfei Wang, James R Torpy, Nenad Bartonicek, et al. A single-cell and spatially resolved atlas of human breast cancers. Nature genetics, 53(9):1334– 1347, 2021.

[22] Quy H Nguyen, Nicholas Pervolarakis, Kerrigan Blake, Dennis Ma, Ryan Tevia Davis, Nathan James, Anh T Phung, Elizabeth Willey, Raj Kumar, Eric Jabart, et al. Profiling human breast epithelial cells using single cell rna sequencing identifies cell diversity. Nature communications, 9(1):2028, 2018.

[23] Mikhail Binnewies, Edward W Roberts, Kelly Kersten, Vincent Chan, Douglas F Fearon, Miriam Merad, Lisa M Coussens, Dmitry I Gabrilovich, Suzanne Ostrand-Rosenberg, Catherine C Hedrick, et al. Understanding the tumor immune microenvironment (time) for effective therapy. Nature medicine, 24(5):541–550, 2018.

[24] Dmitriy Kedrin, Jacco van Rheenen, Lorena Hernandez, John Condeelis, and Jeffrey E Segall. Cell motility and cytoskeletal regulation in invasion and metastasis. Journal of mammary gland biology and neoplasia, 12:143–152, 2007.

[25] Paul J Van Diest, Elsken van der Wall, and Jan PA Baak. Prognostic value of proliferation in invasive breast cancer: a review. Journal of clinical pathology, 57(7):675–681, 2004.

[26] Aravind Subramanian, Pablo Tamayo, Vamsi K Mootha, Sayan Mukherjee, Benjamin L Ebert, Michael A Gillette, Amanda Paulovich, Scott L Pomeroy, Todd R Golub, Eric S Lander, et al. Gene set enrichment analysis: a knowledge-based approach for interpreting genome-wide expression profiles. Proceedings of the National Academy of Sciences, 102(43):15545–15550, 2005.

[27] Sandra Tietscher, Johanna Wagner, Tobias Anzeneder, Claus Langwieder, Martin Rees, Bettina Sobottka, Natalie de Souza, and Bernd Bodenmiller. A comprehensive single-cell map of t cell exhaustion-associated immune environments in human breast cancer. Nature Communications, 14(1):98, 2023.

[28] Talmadge E King, Annie Pardo, and Moiśes Selman. Idiopathic pulmonary fibrosis. The Lancet, 378(9807):1949–1961, 2011.

[29] Jan M Van Deursen. The role of senescent cells in ageing. Nature, 509(7501):439–446, 2014.

[30] Marissa J Schafer, Thomas A White, Koji Iijima, Andrew J Haak, Giovanni Ligresti, Elizabeth J Atkinson, Ann L Oberg, Jodie Birch, Hanna Salmonowicz, YI Zhu, et al. Cellular senescence mediates fibrotic pulmonary disease. Nature communications, 8(1):14532, 2017.

[31] Jodie Birch and Jesús Gil. Senescence and the sasp: many therapeutic avenues. Genes & development, 34(23-24):1565–1576, 2020.

[32] SenNet Consortium. Nih sennet consortium to map senescent cells throughout the human lifespan to understand physiological health. Nature aging, 2(12):1090–1100, 2022.

[33] Dominik Saul, Robyn Laura Kosinsky, Elizabeth J Atkinson, Madison L Doolittle, Xu Zhang, Nathan K LeBrasseur, Robert J Pignolo, Paul D Robbins, Laura J Niedernhofer, Yuji Ikeno, et al. A new gene set identifies senescent cells and predicts senescence-associated pathways across tissues. Nature communications, 13(1):4827, 2022.

[34] AL Fridman and MA Tainsky. Critical pathways in cellular senescence and immortalization revealed by gene expression profiling. Oncogene, 27(46):5975–5987, 2008.

[35] Roberto A Avelar, Javier Gómez Ortega, Robi Tacutu, Eleanor J Tyler, Dominic Bennett, Paolo Binetti, Arie Budovsky, Kasit Chatsirisupachai, Emily Johnson, Alex Murray, et al. A multidimensional systems biology analysis of cellular senescence in aging and disease. Genome biology, 21(1):1–22, 2020.

[36] Aditi U Gurkar, Akos A Gerencser, Ana L Mora, Andrew C Nelson, Anru R Zhang, Anthony B Lagnado, Archibald Enninful, Christopher Benz, David Furman, Delphine Beaulieu, et al. Spatial mapping of cellular senescence: emerging challenges and opportunities. Nature aging, 3(7):776–790, 2023.

[37] Yee Voan Teo, Nattaphong Rattanavirotkul, Nelly Olova, Angela Salzano, Andrea Quintanilla, Nuria Tarrats, Christos Kiourtis, Miryam Müller, Anthony R Green, Peter D Adams, et al. Notch signaling mediates secondary senescence. Cell Reports, 27(4):997–1007, 2019.

[38] Tesfahun Dessale Admasu, Michael J Rae, and Alexandra Stolzing. Dissecting primary and secondary senescence to enable new senotherapeutic strategies. Ageing Research Reviews, 70:101412, 2021.

[39] Matthew Hoare, Yoko Ito, Tae-Won Kang, Michael P Weekes, Nicholas J Matheson, Daniel A Patten, Shishir Shetty, Aled J Parry, Suraj Menon, Rafik Salama, et al. Notch1 mediates a switch between two distinct secretomes during senescence. Nature cell biology, 18(9):979–992, 2016.

[40] Shane A Evans, Yee Voan Teo, Kelly Clark, Takahiro Ito, John M Sedivy, and Nicola Neretti. Single cell transcriptomics reveals global markers of transcriptional diversity across different forms of cellular senescence. bioRxiv, pages 2021–06, 2021.

[41] Elo Madissoon, Amanda J Oliver, Vitalii Kleshchevnikov, Anna Wilbrey-Clark, Krzysztof Polanski, Nathan Richoz, Ana Ribeiro Orsi, Lira Mamanova, Liam Bolt, Rasa Elmentaite, et al. A spatially resolved atlas of the human lung characterizes a gland-associated immune niche. Nature Genetics, 55(1):66–77, 2023.

[42] Gene Ontology Consortium. The gene ontology (go) database and informatics resource. Nucleic acids research, 32(suppl 1):D258–D261, 2004.

[43] Bennett G Childs, Martina Gluscevic, Darren J Baker, Remi-Martin Laberge, Dan Marquess, Jamie Dananberg, and Jan M Van Deursen. Senescent cells: an emerging target for diseases of ageing. Nature reviews Drug discovery, 16(10):718–735, 2017.

[44] Audrey Lasry and Yinon Ben-Neriah. Senescence-associated inflammatory responses: aging and cancer perspectives. Trends in immunology, 36(4):217–228, 2015.

[45] Jian Hu, Xiangjie Li, Kyle Coleman, Amelia Schroeder, Nan Ma, David J Irwin, Edward B Lee, Russell T Shinohara, and Mingyao Li. Spagcn: Integrating gene expression, spatial location and histology to identify spatial domains and spatially variable genes by graph convolutional network. Nature methods, 18(11):1342–1351, 2021.

[46] Kangning Dong and Shihua Zhang. Deciphering spatial domains from spatially resolved transcriptomics with an adaptive graph attention auto-encoder. Nature communications, 13(1):1739, 2022.

[47] Wei-Ting Chen, Ashley Lu, Katleen Craessaerts, Benjamin Pavie, Carlo Sala Frigerio, Nikky Corthout, Xiaoyan Qian, Jana Laláková, Malte Kühnemund, Iryna Voytyuk, et al. Spatial transcriptomics and in situ sequencing to study alzheimer’s disease. Cell, 182(4):976–991, 2020.

[48] Anna Lyubetskaya, Brian Rabe, Andrew Fisher, Anne Lewin, Isaac Neuhaus, Constance Brett, Todd Brett, Ethel Pereira, Ryan Golhar, Sami Kebede, et al. Assessment of spatial transcriptomics for oncology discovery. Cell Reports Methods, 2(11), 2022.

## References

[1] Ludvig Bergenstråhle, Bryan He, Joseph Bergenstråhle, Xesús Abalo, Reza Mirzazadeh, Kim Thrane, Andrew L Ji, Alma Andersson, Ludvig Larsson, Nathalie Stakenborg, et al. Super-resolved spatial transcriptomics by deep data fusion. Nature biotechnology, 40(4):476–479, 2022.

[2] Hao Chen, Dongshunyi Li, and Ziv Bar-Joseph. Scs: cell segmentation for high-resolution spatial transcriptomics. Nature methods, 20:1237–1243, 2023.

[3] Serge Beucher. Use of watersheds in contour detection. In Proc. Int. Workshop on Image Processing, *Sept.* 1979, pages 17–21, 1979.

[4] Amit Zeisel, Hannah Hochgerner, Peter Lönnerberg, Anna Johnsson, Fatima Memic, Job Van Der Zwan, Martin Häring, Emelie Braun, Lars E Borm, Gioele La Manno, et al. Molecular architecture of the mouse nervous system. Cell, 174(4):999–1014, 2018.

[5] Alma Andersson, Joseph Bergenstråhle, Michaela Asp, Ludvig Bergenstråhle, Aleksandra Jurek, Jośe Ferńandez Navarro, and Joakim Lundeberg. Single-cell and spatial transcriptomics enables probabilistic inference of cell type topography. Communications biology, 3(1):565, 2020.

[6] Vitalii Kleshchevnikov, Artem Shmatko, Emma Dann, Alexander Aivazidis, Hamish W King, Tong Li, Rasa Elmentaite, Artem Lomakin, Veronika Kedlian, Adam Gayoso, et al. Cell2location maps fine-grained cell types in spatial transcriptomics. Nature biotechnology, 40(5):661–671, 2022.

[7] Ying Ma and Xiang Zhou. Spatially informed cell-type deconvolution for spatial transcriptomics. Nature biotechnology, 40(9):1349–1359, 2022.

[8] Patrik L Ståhl, Fredrik Salmén, Sanja Vickovic, Anna Lundmark, Jośe Ferńandez Navarro, Jens Magnusson, Stefania Giacomello, Michaela Asp, Jakub O Westholm, Mikael Huss, et al. Visualization and analysis of gene expression in tissue sections by spatial transcriptomics. Science, 353(6294):78–82, 2016.

[9] SenNet Consortium. Nih sennet consortium to map senescent cells throughout the human lifespan to understand physiological health. Nature aging, 2(12):1090–1100, 2022.

[10] Nam D Nguyen, Lorena Rosas, Timur Khaliullin, Peiran Jiang, Euxhen Hasanaj, Jose A Ovando, Marta Bueno, Melanie Konigshoff, Oliver Eickelberg, Mauricio Rojas, et al. Optimal transport for mapping senescent cells in spatial transcriptomics. bioRxiv, pages 2023–08, 2023.

[11] Sunny Z Wu, Ghamdan Al-Eryani, Daniel Lee Roden, Simon Junankar, Kate Harvey, Alma Andersson, Aatish Thennavan, Chenfei Wang, James R Torpy, Nenad Bartonicek, et al. A single-cell and spatially resolved atlas of human breast cancers. Nature genetics, 53(9):1334– 1347, 2021.

[12] F Alexander Wolf, Philipp Angerer, and Fabian J Theis. Scanpy: large-scale single-cell gene expression data analysis. Genome biology, 19:1–5, 2018.

[13] Lisa Sikkema, Ciro Ramírez-Suástegui, Daniel C Strobl, Tessa E Gillett, Luke Zappia, Elo Madissoon, Nikolay S Markov, Laure-Emmanuelle Zaragosi, Yuge Ji, Meshal Ansari, et al. An integrated cell atlas of the lung in health and disease. Nature Medicine, pages 1–15, 2023.

[14] Mohammad Lotfollahi, Mohsen Naghipourfar, Malte D Luecken, Matin Khajavi, Maren Büttner, Marco Wagenstetter, Žiga Avsec, Adam Gayoso, Nir Yosef, Marta Interlandi, et al. Mapping single-cell data to reference atlases by transfer learning. Nature biotechnology, 40(1):121–130, 2022.

[15] Matthew Hoare, Yoko Ito, Tae-Won Kang, Michael P Weekes, Nicholas J Matheson, Daniel A Patten, Shishir Shetty, Aled J Parry, Suraj Menon, Rafik Salama, et al. Notch1 mediates a switch between two distinct secretomes during senescence. Nature cell biology, 18(9):979–992, 2016.

[16] Shane A Evans, Yee Voan Teo, Kelly Clark, Takahiro Ito, John M Sedivy, and Nicola Neretti. Single cell transcriptomics reveals global markers of transcriptional diversity across different forms of cellular senescence. bioRxiv, pages 2021–06, 2021.

[17] Yee Voan Teo, Nattaphong Rattanavirotkul, Nelly Olova, Angela Salzano, Andrea Quintanilla, Nuria Tarrats, Christos Kiourtis, Miryam Müller, Anthony R Green, Peter D Adams, et al. Notch signaling mediates secondary senescence. Cell Reports, 27(4):997–1007, 2019.

