## Supplementary Figures and Tables for "scResolve: Recovering single cell expression profiles from multi-cellular spatial transcriptomics"

* These authors contributed equally.

### Supplentary Figures

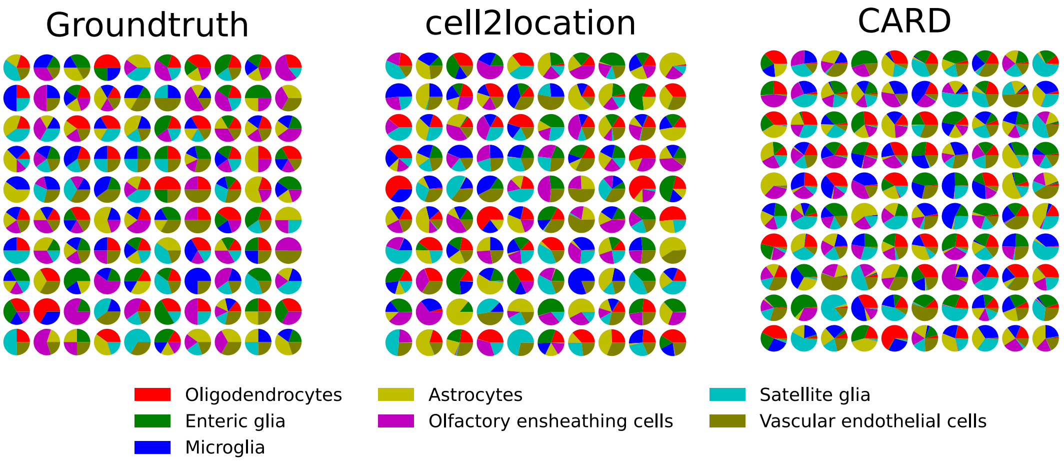


Supplementary Figure 1: . **Cell type composition from ground truth data and cell type deconvolution methods**. Left, Ground truth cell type composition of each simulated spot in Figure 2a. Middle, Reconstructed cell type compositions from cell2location. Right, Reconstructed cell type compositions from CARD.

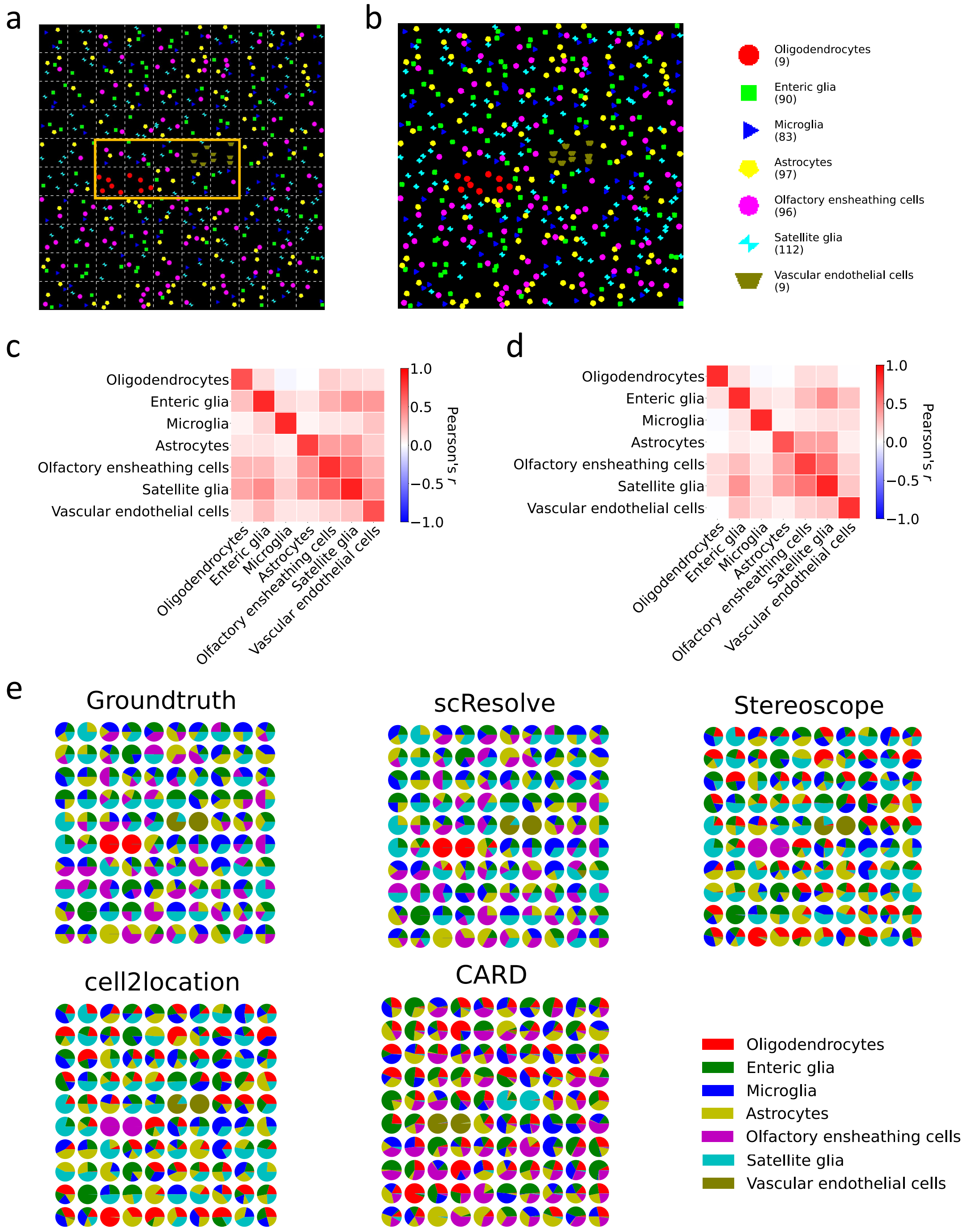


Supplementary Figure 2: **Simulation study for rare cell type.** **a**, Image for the synthetic data section with simulated rare cell types, where Oligodendrocytes (red circle) and Vascular endothelial cells (brown trapezoid) were used as rare cell types. The whole spatial map was divided into a grid to simulate multicellular spots. The yellow box indicates the region surrounding the cells of rare cell types. **b**, Recovered single cells with their annotated cell types in the spatial map. **c**, Expression correlation between cell types in scResolve recovered cells and those in real cells for sinlge cells in **a-b**. **d**, Expression correlation between cell types in real cells. **e**, Cell type composition of spots in **a** computed from the ground truth data and reconstructed from different methods.

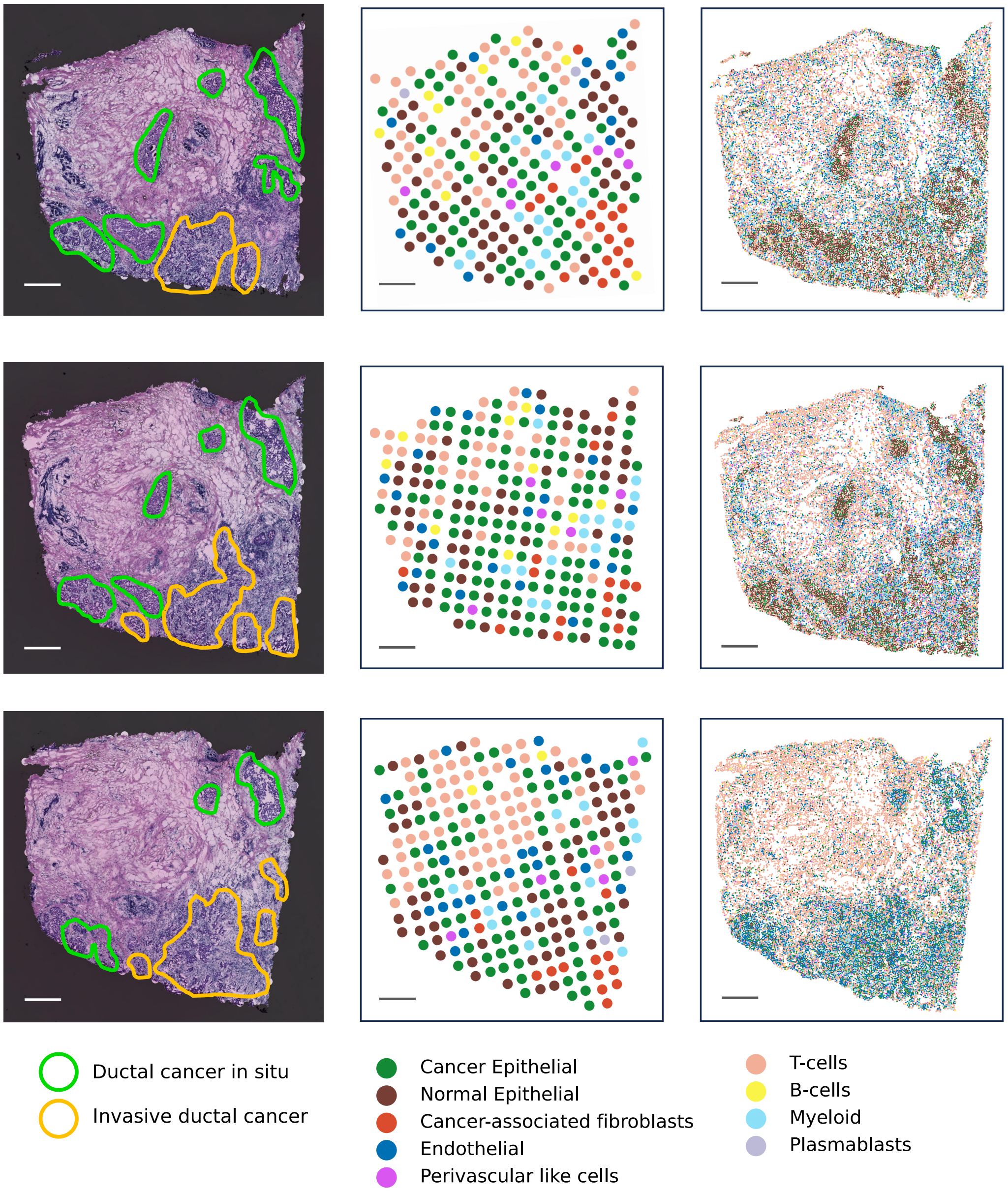


Supplementary Figure 3: **Results for the three other sections in the breast cancer dataset**. Left, Histology images of the three tissue sections. Middle, Cell type annotations on the original ST spots. Right, Cell type annotations on the scResolve recovered single cells. Scale bars, 500 $\mu$m.

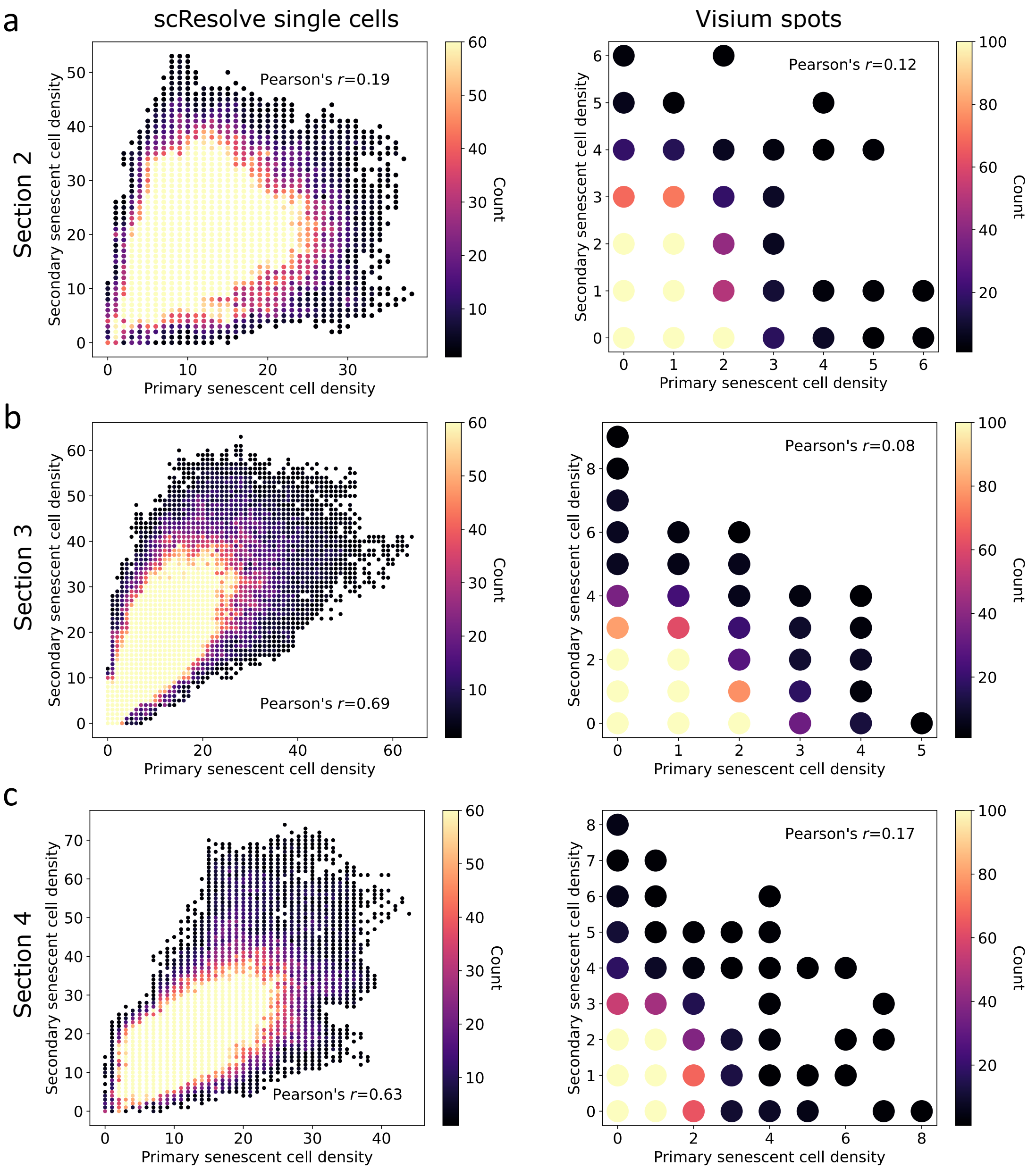


Supplementary Figure 4: **Spatial correlation between primary and secondary senescent cells**. **a-c**, Results are shown for the top, middle, and bottom sections in Figure 4l. Left, Correlation between the densities of primary and secondary senescent cells at the single cell level. Right, Correlation between the densities of primary and secondary senescent cells and at the spot level.

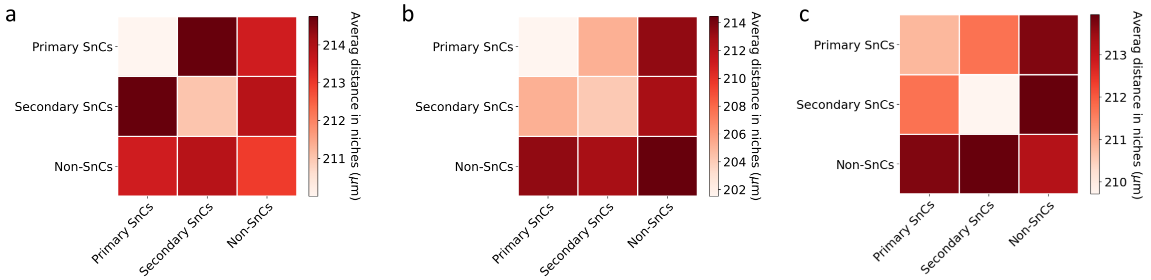


Supplementary Figure 5: **Heatmap of the average distance between cells of different cell populations in niches**. **a-c**, Results are shown for the top, middle, and bottom sections in Figure 4l.

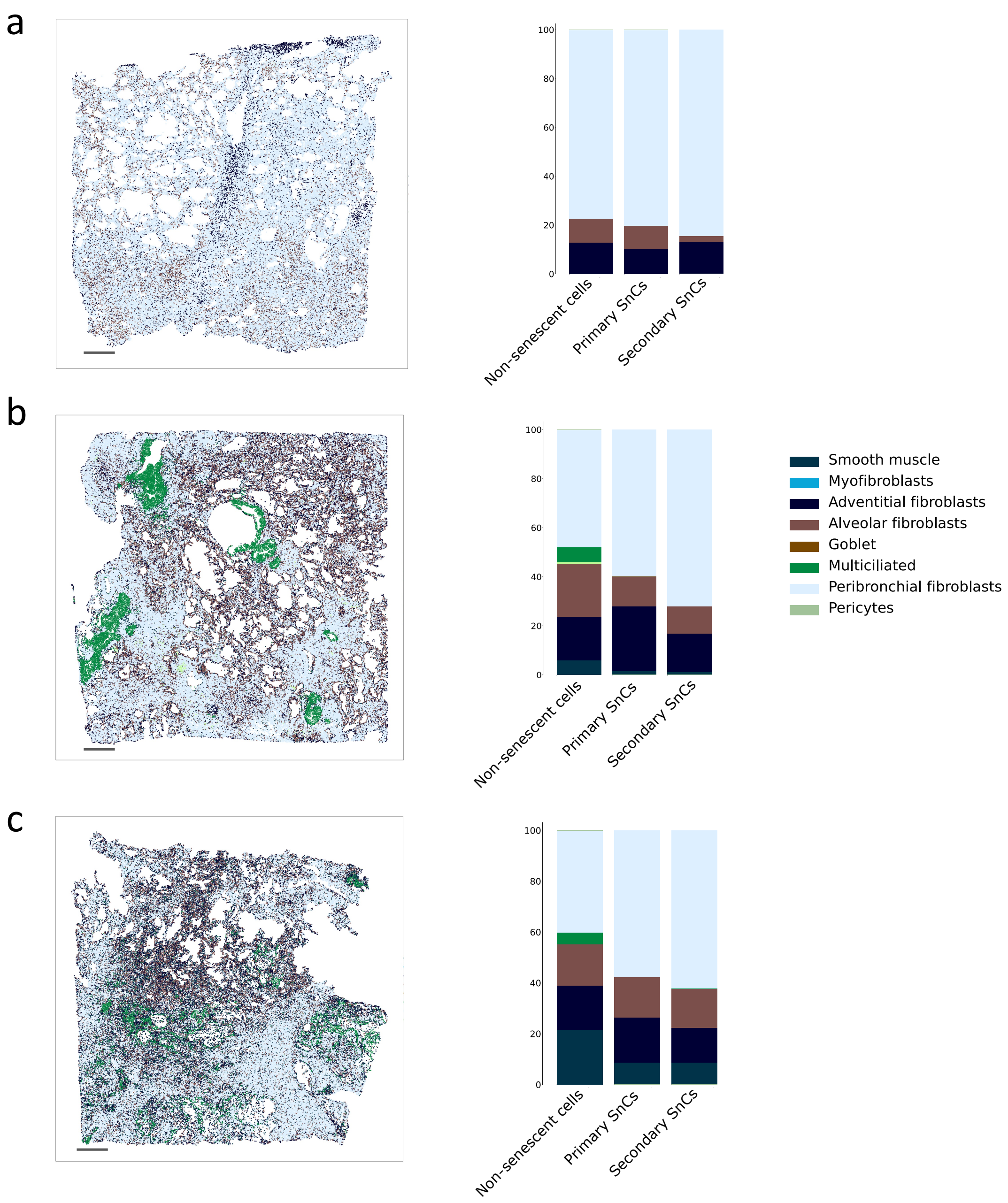


Supplementary Figure 6: **Cell type annotations and cell type compositions for the other three IPF sections**. **a-c**, Results are shown for the top, middle, and bottom sections in Figure 4l. Left, Cell type annotation of recovered single cells from scResolve. Right, Cell type compositions for different groups of cells. Scale bars, 500 $\mu$m.

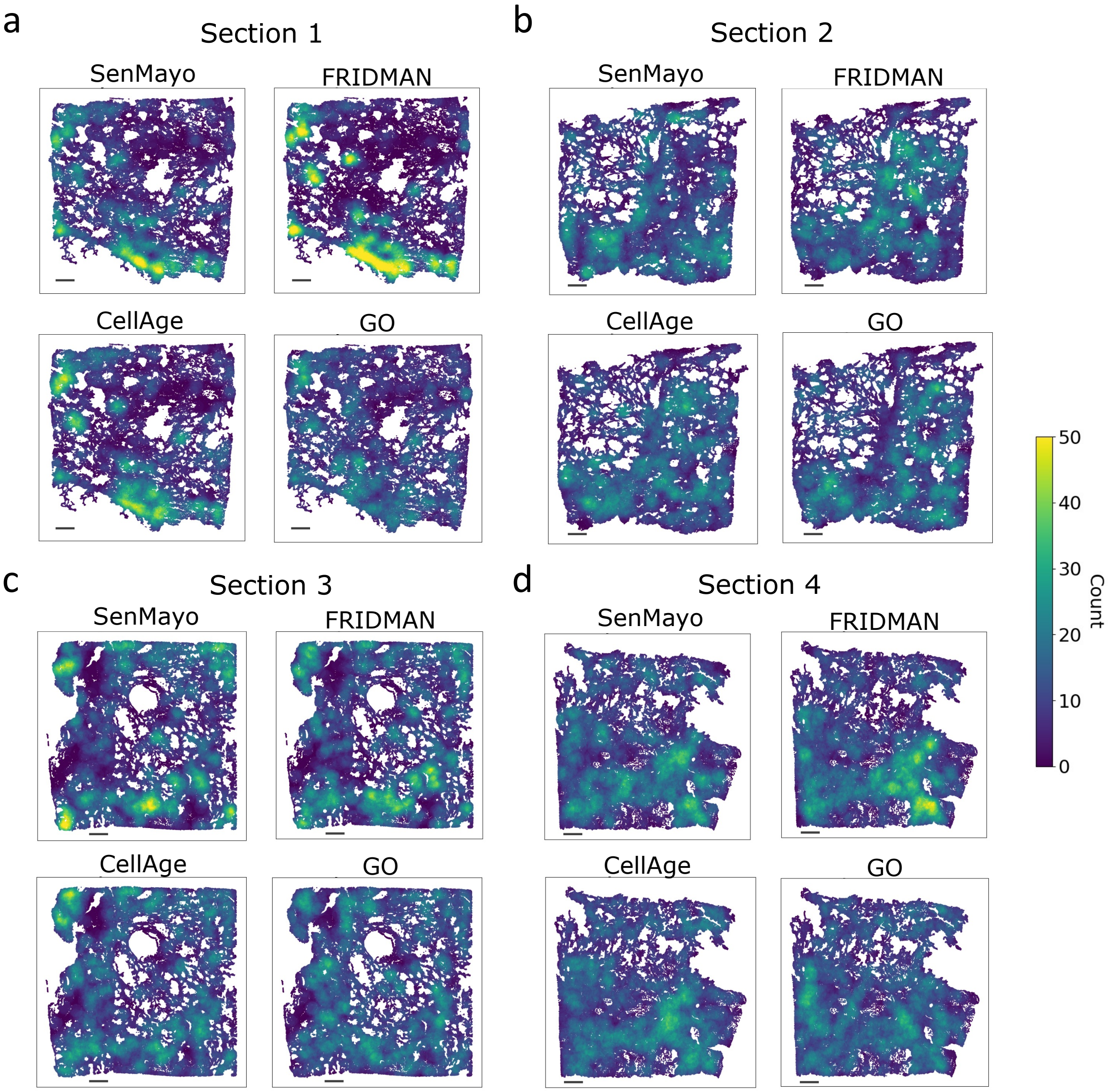


Supplementary Figure 7: **Senescent cell densities in different IPF sections**. **a-d**, The density of senescent cells in four different sections. For each section, four different gene lists (SenMayo, FRIDMAN, CellAge, and GO) were used as markers for identifying senescent cells. Scale bars, 500 $\mu$m.

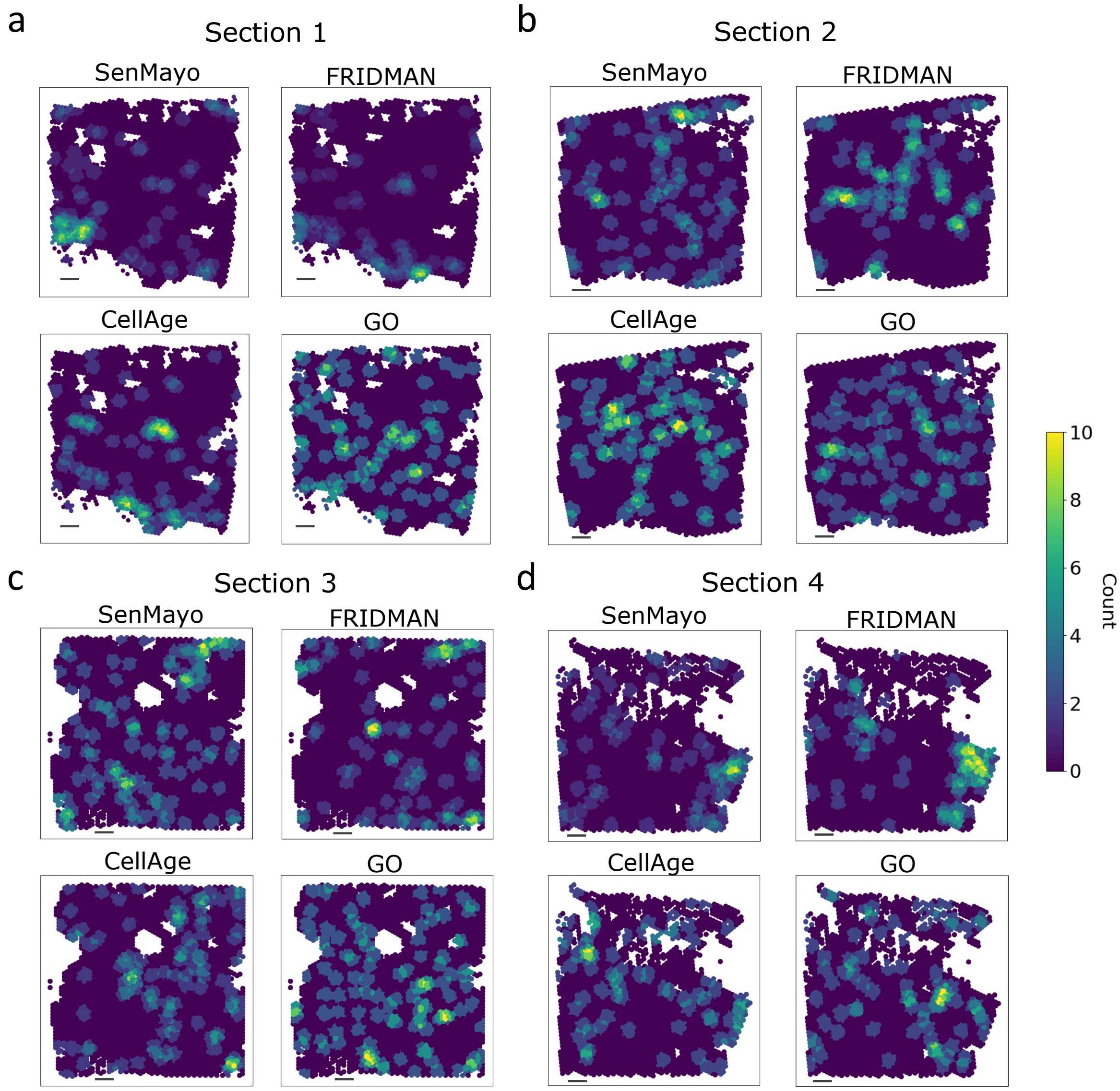


Supplementary Figure 8: **Senescent cell densities in different IPF sections at the spot level**. **a-d**, The density of senescent cells in four different sections. For each section, four different gene lists (SenMayo, FRIDMAN, CellAge, and GO) were used as markers for identifying spots that contain senescent cells. Scale bars, 500 $\mu$m.

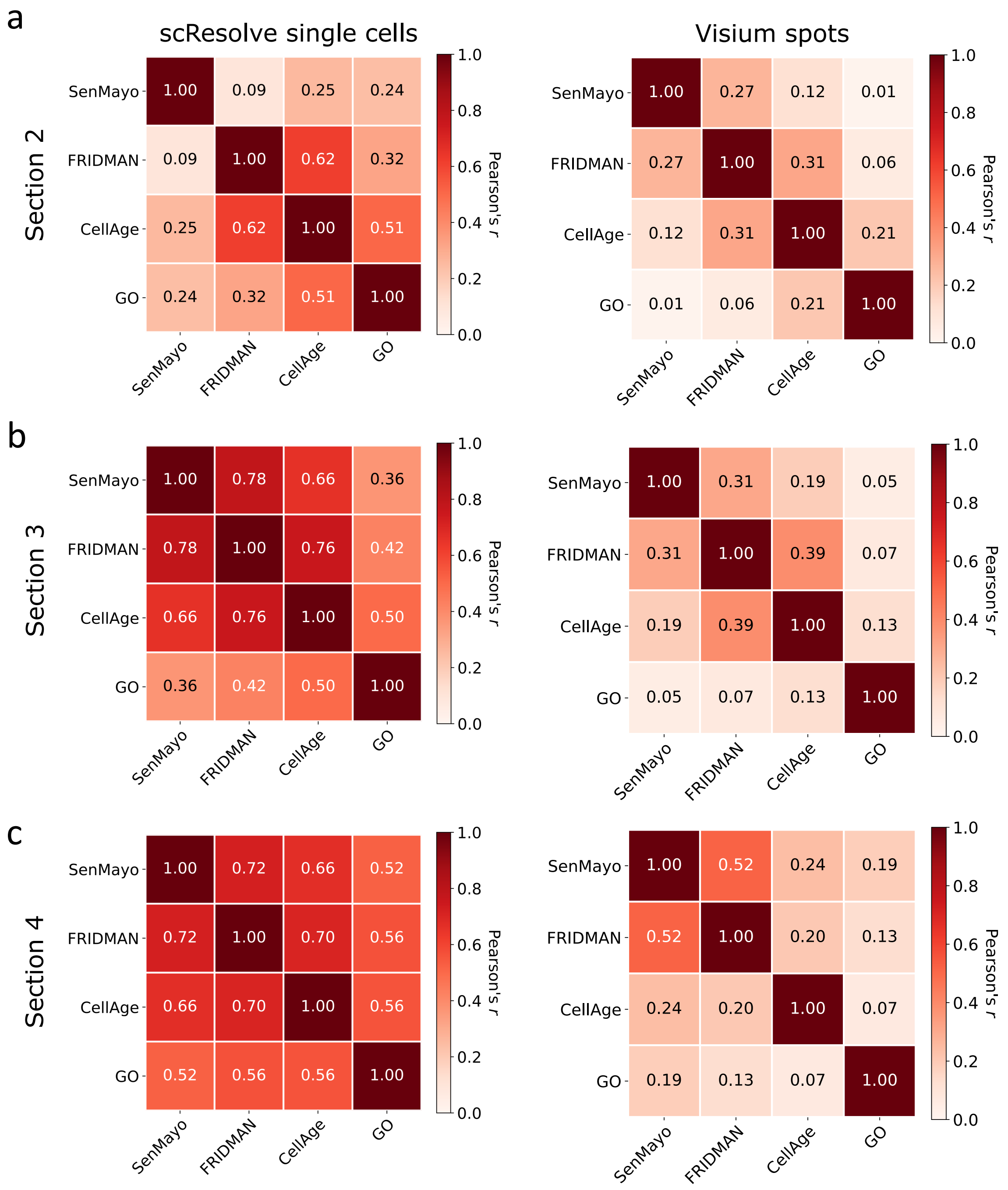


Supplementary Figure 9: **Consistency between different gene lists in identifying senescent cells**. **a-c**, Results are shown for the top, middle, and bottom sections in Figure 4l. Left: The spatial correlation among the densities of senescent cells identified using different marker gene lists. Right: The spatial correlation among the densities of spots containing senescent cells identified using different marker gene lists.

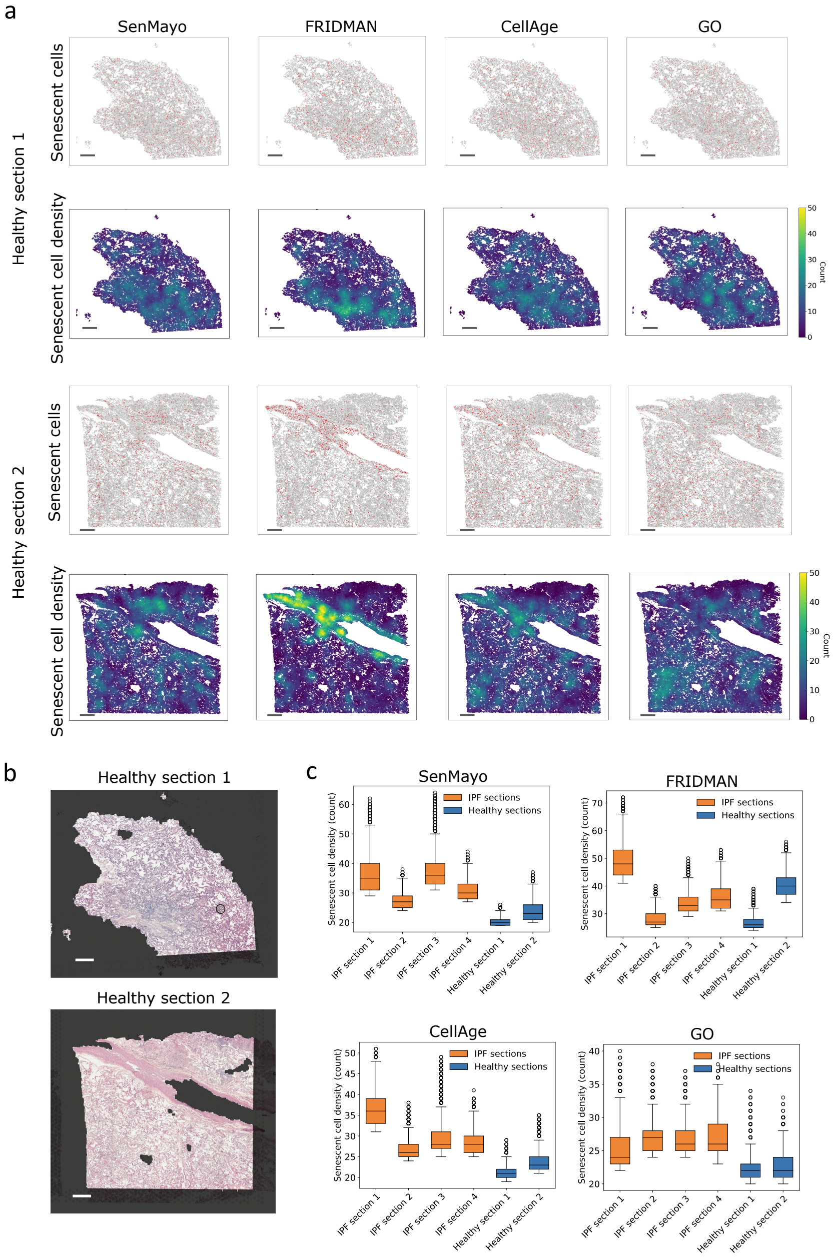


Supplementary Figure 10: **Application of scResolve to two healthy lung sections**. **a**, Identified senescent cells (top) and senescent cell density around each cell location (bottom) in the recovered single cells. Results are shown for senescent cells identified using different gene lists. **b**, The histology images of the two healthy lung sections. **c**, Densities of senescent cells in the top 10% cell locations with the highest senescent cell densities in each section. IPF tissue sections in general contain more locations with higher senescent cell densities than healthy tissue sections across all four gene lists. The only exception is for the FRIDMAN gene list in healthy section 2, where the identified senescent cells show high density in specific regions. However, the regions correspond to the Bronchiolo-vascular bundles with associated interstitium, where smooth muscle cells and blood vessel-related cells are enriched [1]. Genes associated with blood vessel development (GO:0001568) and regulation of smooth muscle cell proliferation (GO:0048660) were found to be enriched in the FRIDMAN gene list [2], which explains why the FRIDMAN gene list identifies more cells in these regions. Scale bars, 500 $\mu$m.

a

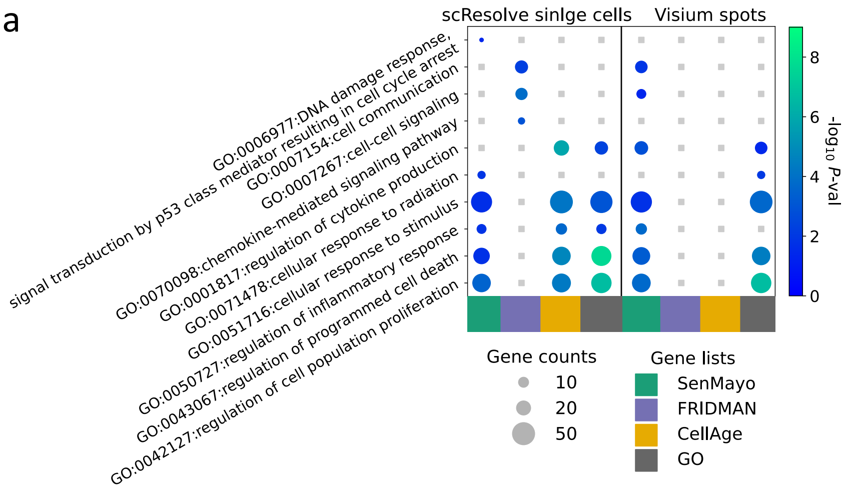


b

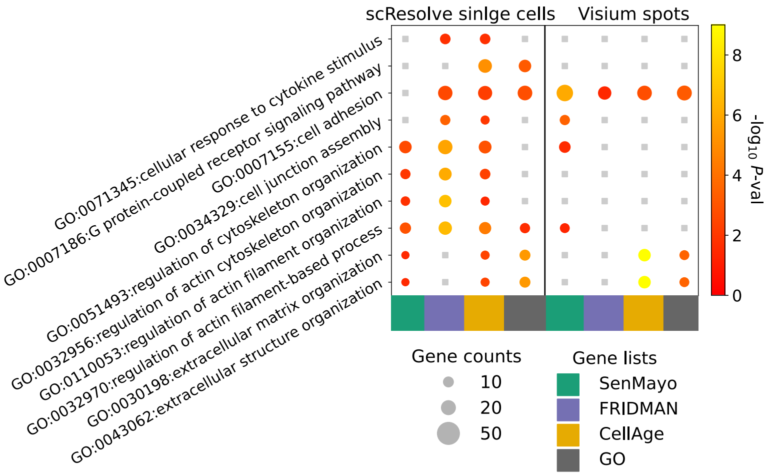


Supplementary Figure 11: **Differential expression analysis between primary and secondary senescent cells**. **a**, Enrichment analysis of Gene Ontology (GO) in the top 300 differentially expressed genes after removing marker genes of primary senescent cells. **b**, GO enrichment analysis in the top 300 differentially expressed genes after removing marker genes of secondary senescent cells. The size of the dots indicates the number of genes associated with the GO terms present in the differentially expressed genes, while the color of the dots reflects the corresponding $P$-values.

### Supplentary Tables

**Supplementary Table 1: Benchmarks of performance.** The Pearson Correlation coefficient (PCC) and Mean Squared Error (MSE) were calculated between groundtruth cell type composition and predicted cell type composition. The average values and standard deviations for five replicates are shown for each method and section.

| | PCC | | MSE | |
|---|---|---|---|---|
| | Uniform | Rare | Uniform | Rare |
| Stereoscope | 0.46(0.0000) | 0.47(0.0000) | 0.85(0.0000) | 1.50(0.0000) |
| cell2location | 0.46(0.0005) | 0.47(0.0009) | 0.89(0.0002) | 1.51(0.0013) |
| Card | 0.01(0.0000) | 0.28(0.0000) | 0.93(0.0000) | 1.69(0.0000) |
| scResolve | **0.90**(0.1019) | **0.96**(0.0159) | **0.14**(0.1414) | **0.14**(0.0167) |

### References

[1] Teri J Franks et al. "Resident cellular components of the human lung: current knowledge and goals for research on cell phenotyping and function". In: *Proceedings of the American thoracic society* 5.7 (2008), pp. 763–766.

[2] Yingyao Zhou et al. "Metascape provides a biologist-oriented resource for the analysis of systems-level datasets". In: *Nature communications* 10.1 (2019), p. 1523.
